# Multiplexed imaging of G-proteins and ERK activity upon activation of CaSR

**DOI:** 10.1101/2025.11.19.689196

**Authors:** Sergei Chavez-Abiega, Luca Bordes, Marten Postma, Frank J. Bruggeman, J. Goedhart

## Abstract

We have previously shown that the calcium sensing receptor (CaSR) activates different G proteins and second messengers in single cells, and that GPCR and G protein-mediated ERK activity can be highly dynamic and heterogeneous at the single cell level. Here, we attempted to investigate the CaSR-Gi-ERK signaling pathway in single cells with previously characterized biosensors. We developed a strategy to simultaneously track G-protein and ERK activities in a single cell, using FRET-based and translocation sensors, a novel large-Stokes-shift green fluorescent protein, and an imaging setup with a single detector and a single wavelength for excitation. Extensive characterization and optimization to unmix the signals showed that the differences in dynamic range and type of read-out obtained from these biosensors made it very challenging to obtain robust FRET-measurements in the developed setup. When focusing solely on ERK, we found signs of possible ERK activity in response to calcium stimulation via CaSR, as well as milder changes in the absence of calcium treatment. Further optimization of the model, probes, and image processing will be necessary to develop a setup to robustly measure FRET and translocation simultaneously in single cells. We present the challenges we encountered and discuss future developments that can overcome them.

## INTRODUCTION

G-protein coupled receptor (GPCR) signaling networks are very complex and compose several pathways that have unique spatial and temporal dynamics. Live cell imaging techniques with fluorescence-based biosensors are instrumental to study these processes, as they allow us to look into biological events in real time in individual cells [1]. There exists however a limited number of fluorophores that can be used simultaneously due to overlap in the excitation and emission spectra. By using the well-chosen combinations of biosensors and fluorophores it is possible to image up to 6 biosensors simultaneously [2].

The first mediators of the signaling pathways activated by the GPCRs are the heterotrimeric G-proteins present at the plasma membrane. G-proteins can activate ERK through different pathways, and it is now known that β-arrestins can also activate specific ERK pools. However, it is not clear whether ERK activation by β-arrestins is a process that occurs independently of G-proteins, since two recent studies suggest that G proteins are essential [3,4]. Despite the controversy, most studies that study G-protein vs β-arrestin ligand-biased signaling use ERK activity as the read-out for β-arrestin.

One of the most studied GPCRs in ligand-biased studies is the calcium sensing receptor (CaSR). The reasons are that activity of this class C GPCR is modulated by different chemical classes of molecules and that it couples to activate multiple G-protein families [5]. Here we followed G-protein and ERK activities simultaneously in single cells upon modulation of CaSR activity. Available G-protein biosensors are predominantly FRET-based, with the α subunit tagged with a CFP as a donor, and βγ with a YFP as acceptor, or the other way around [6–9]. Following previous reported results [10], where we observed Gi1 heterotrimer dissociation upon CaSR activation, and the established relationship between Gi and ERK in CaSR signaling [11,12] we decided to use the Gi1 sensor. As for ERK activity, different sensor architectures are available, from FRET-pairs [13–15] to those that employ a single FP or a single channel [2,16,17]. We decided to use a kinase translocation reporter (KTR), that contains a fragment of the transcription factor Elk1, which upon phosphorylation by ERK translocates from the nucleus to the cytoplasm [17]. We showed previously [18] that this KTR has a high dynamic range, and as it allows to choose any FP to follow and quantify the translocation, it offers high versatility when it comes to combining it with other biosensors.

Here, we report a strategy to simultaneously image four probes on a single detector with a single wavelength for excitation. We discuss the choice of probes, the imaging conditions, the image analysis steps, including the unmixing analysis, and the optimization of the sample preparation.

## RESULTS

### Choice of fluorophores

To investigate simultaneously the activities of Gi and ERK in response to CaSR activation, we decided to explore the possibility of using a single excitation wavelength to quantify changes in a FRET-based biosensor and a translocation reporter. For this, we intended to take advantage of an imaging setup that allows to excite the cells with 420/40 nm and obtain four spectrally separated images corresponding to the emissions of CFP, GFP, YFP, and OFP/RFP. A representation of this system is shown in Figure 1.

**Figure 1.**
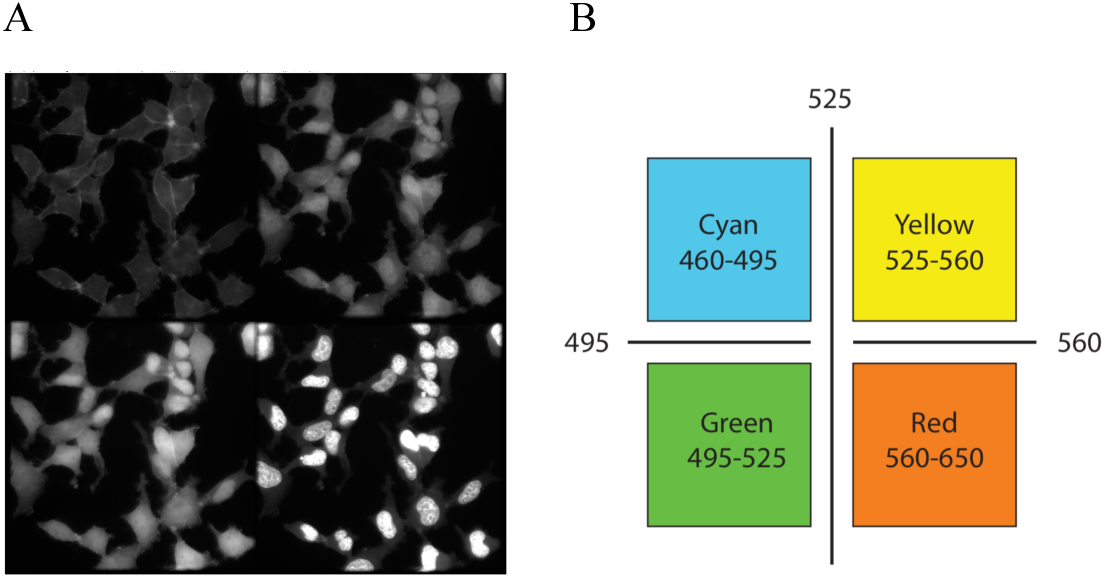
Single-excitation multi-channel system. The sample is illuminated with 400-440 nm, and the emission is first passed through a 460 LP filter and then through three dichroic filters to be split in 4 channels with the following wavelengths: 460-495 nm, 495-525 nm, 525-560 nm, and >560 nm. A: Multi-channel image of cells expressing Gαi1 FRET sensor mTurquoise2-cpVenus, ERK-KTR-lssSGFP2, and H2A-lssmOrange. B: Wavelengths detected in each of the channels.

To complement the CFP-YFP FRET pair and lssSGFP2, we explored published large-Stokes-shift orange and red fluorescent proteins. Based on theoretical brightness from literature and plasmid availability, we selected hmKeima8.5, lssmCherry1, lssmKate2, and lssmOrange. The spectral characteristics of these FPs are listed in Table 1.

**Table 1.**
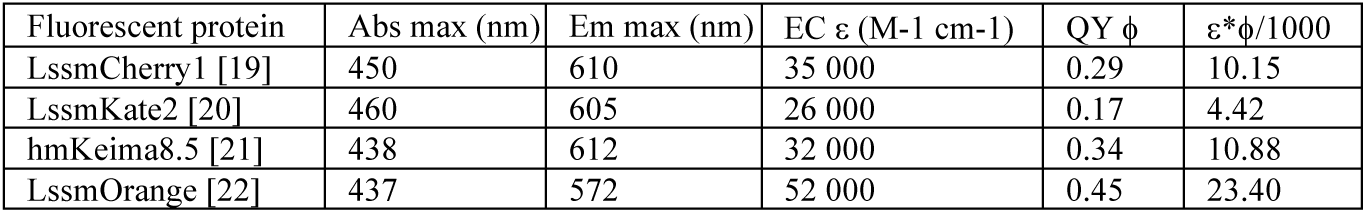
Photophysical properties of the large-Stokes-shift orange and red fluorescent protein candidates. Abs max: Absorbance maximum peak wavelength. Em max: Emission maximum peak wavelength. EC: Extinction coefficient. QY: Quantum yield.

The product of quantum yield and extinction coefficient is often referred to as theoretical brightness, which does not necessarily correlate to the emission observed in cells. To compare the brightness of these candidate fluorescent proteins in cells, we generated tandem constructs containing each candidate FP and mTq2 with a T2A linker in between. This linker results in comparable expression of two fluorescent proteins [23], and it allows to measure the emission of a protein relative to mTq2.

First, we transfected HeLa cells with each of these fusion constructs, and used spectral imaging microscopy to obtain the emission spectrum of each of the fluorophores of interest normalized to the spectrum of mTq2, our reference fluorophore. Figure 2A shows that the area under the curve of the emission spectra of lssmOrange and hmKeima are much larger than those of lssmCherry1 and lssmKate2. Second, we used widefield microscopy to image these transfected cells and quantify the fluorescence emitted by these fluorophores and plotted them against the measured fluorescence of our reference mTq2. As shown in Figure 2B, and in agreement with what we observed using spectral imaging microscopy, lssmOrange and hmKeima8.5 were considerably brighter in live cells compared to lssmCherry and lssmKate2. From these results, we chose lssmOrange as the fourth fluorophore for the project.

**Figure 2.**
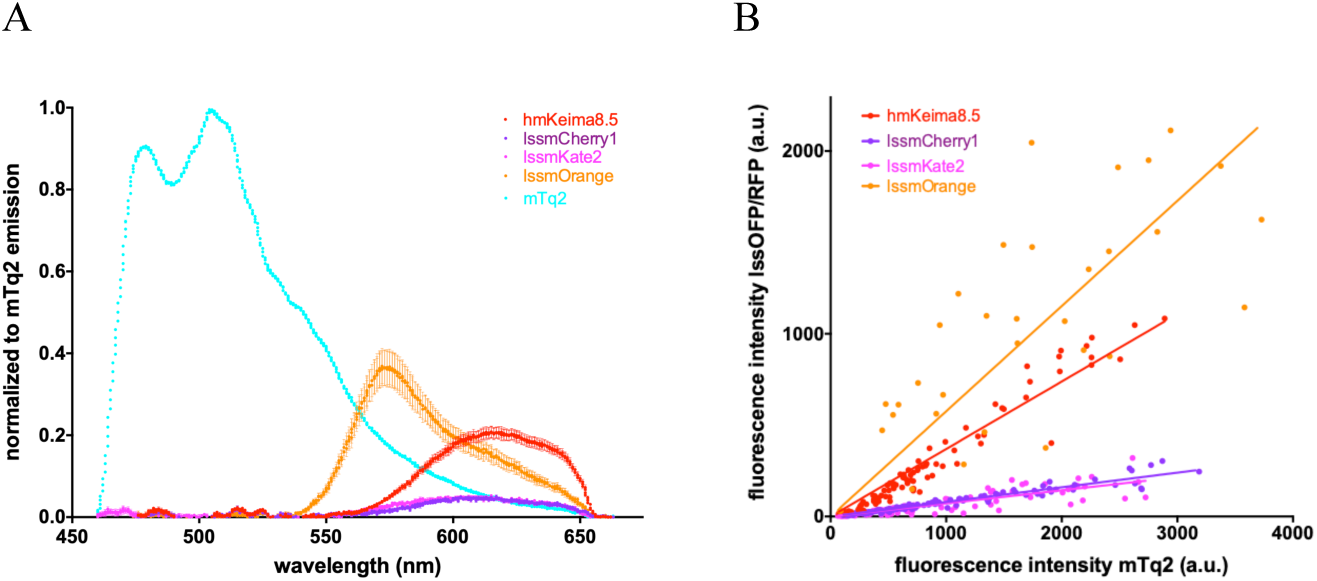
Emission and brightness analysis of candidate lssRFPs. A-B: Live cells were transfected with tandem constructs of each lssRFP and mTq2. A: Emission spectra of hmKeima8.5, lssmCherry1, lssmKate2 and lssmOrange, normalized to the peak of the emission spectrum of mTq2. Spectra were acquired in single cells, and traces show the average and standard deviation of the mean. B: Fluorescence intensity counts in the lssRFP and mTq2 channels are plotted per cell. Each point represents a single cell measurement.

### Unmixing

Despite the fact that these four fluorophores emit light with different colors, a relatively high spectral overlap exists between their emission spectra. This is especially clear for the emission spectrum of mTq2 that overlaps largely with the spectra of lssSGFP2 and SYFP2, as shown in Figure 3. For this reason, it was necessary to unmix the signals and therefore to calculate the bleedthrough percentage per fluorescent protein per channel. We calculated the theoretical bleedthrough using the protein spectra and the microscope hardware information, and we also measured it experimentally in transformed bacteria, transfected HEK293 cells, and purified proteins in Nickel beads. The results are shown in Figure 4 and Table 2.

**Figure 3.**
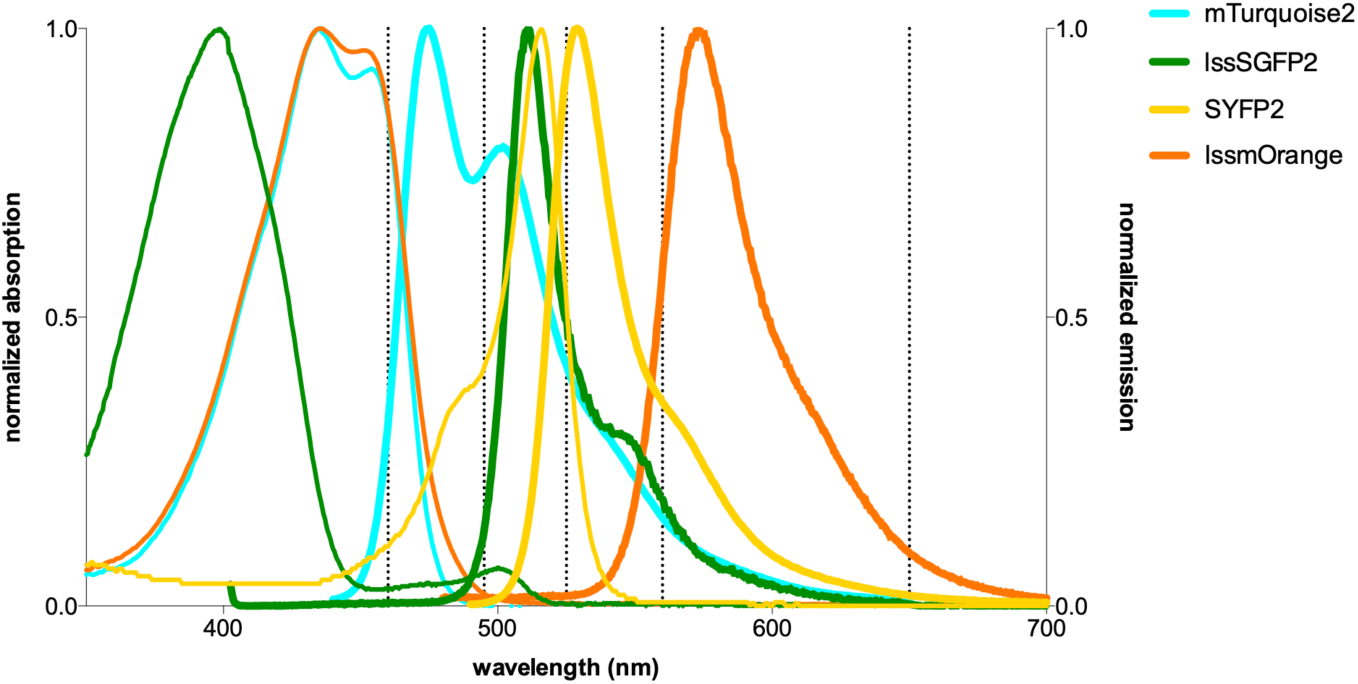
Absorption and emission spectra of the fluorescent proteins used in the multi-channel system. The spectra were acquired by Marieke Mastop [24] and normalized to their peak values. Dashed lines indicate the ranges of wavelengths detected by the four channels in the system.

**Figure 4.**
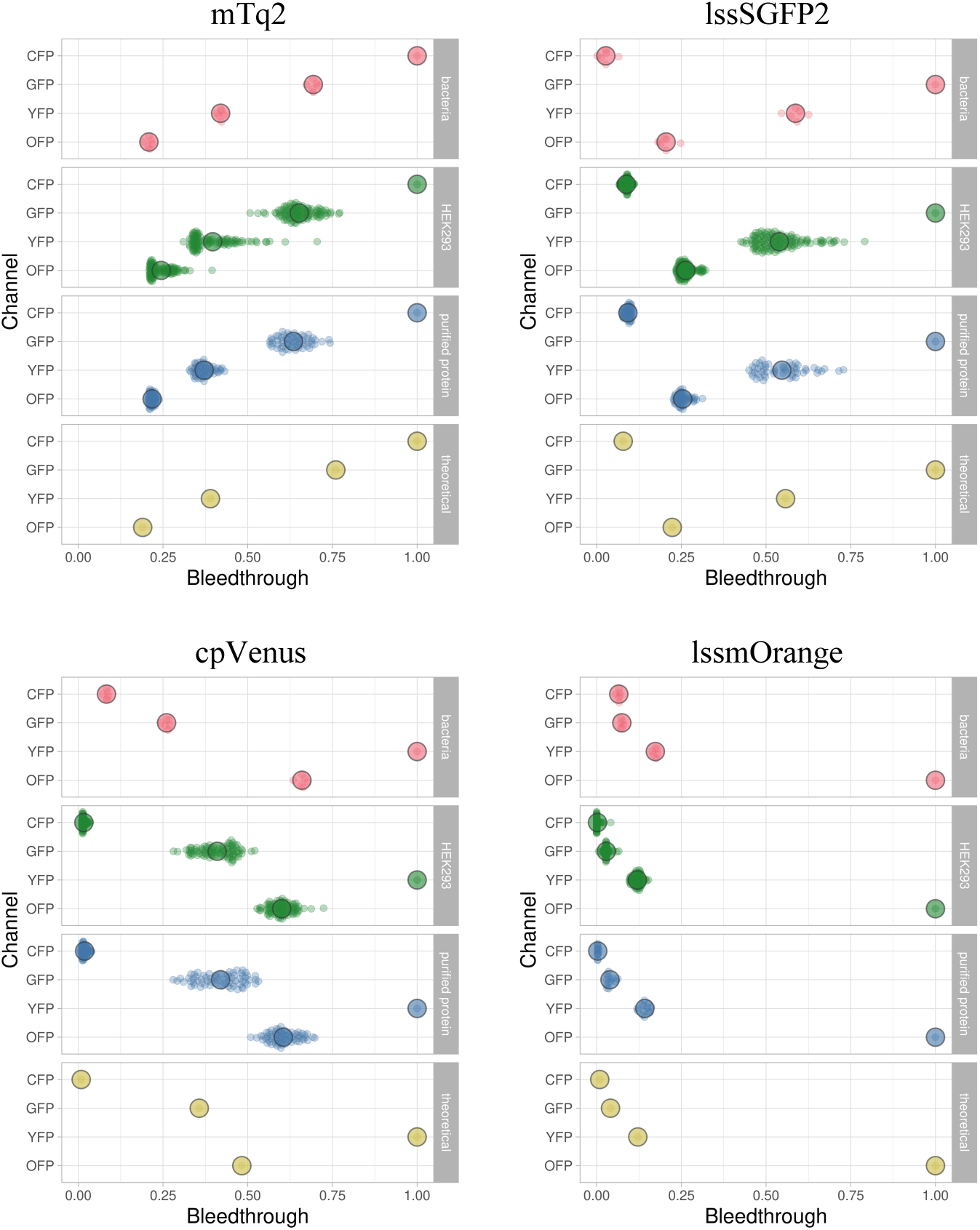
Relative fluorescence of each FP in all four channels calculated by different methods. The bleedthrough is expressed as the fraction per channel relative to the channel with the highest value. It was measured in transformed bacteria, transfected HEK293 cells, purified protein in Nickel beads, and calculated theoretically. Each small dot represents a single intensity measurement: a field of view from a bacterial culture, a single transfected cell, or a single bead. Each large dot represents the average of all single measurements. Data was plotted using SuperPlotsofData [25].

**Table 2.**
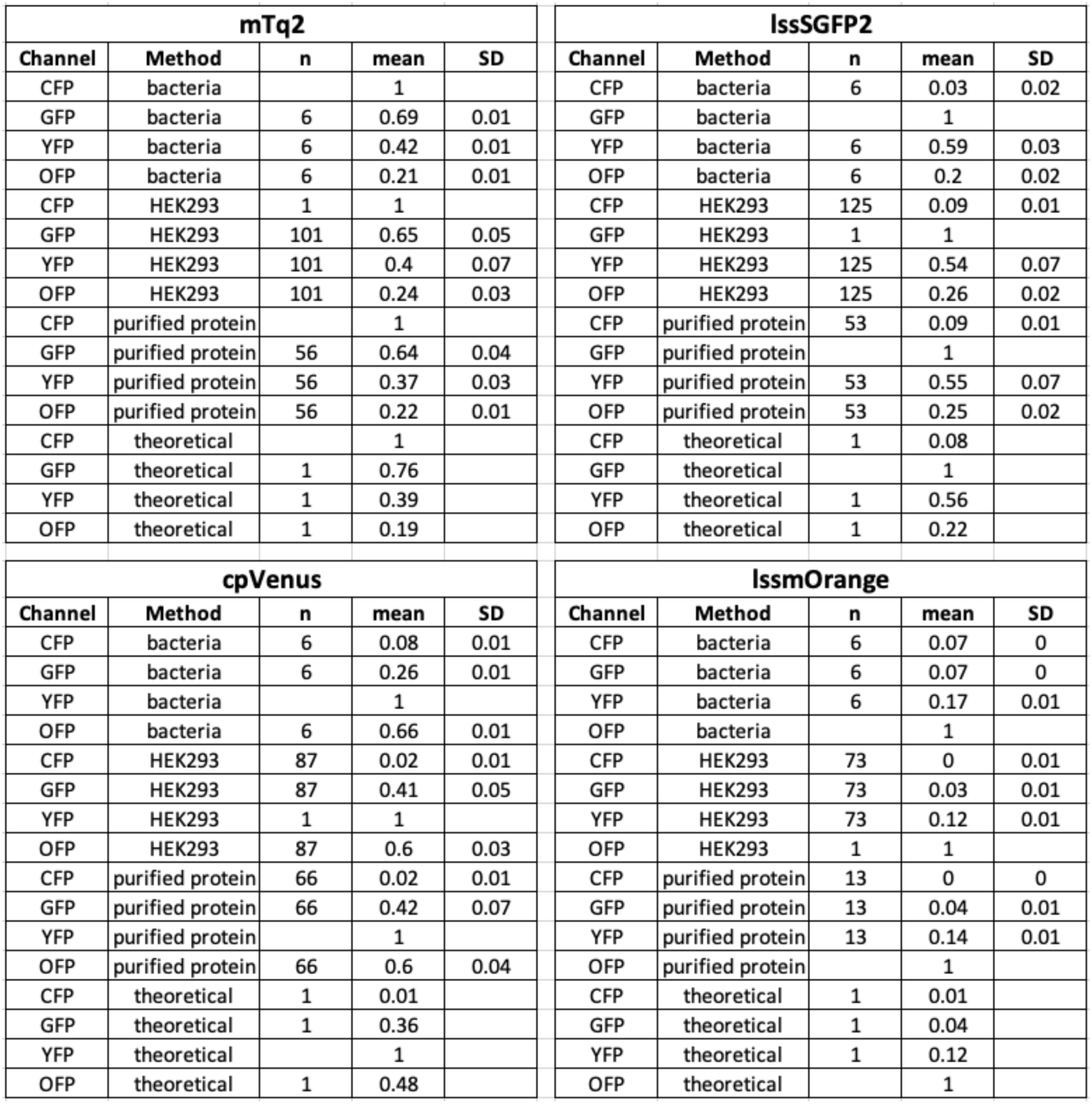
Relative fluorescence of each FP in all four channels calculated by different methods. The average and standard deviation of the measurements per FP per channel were calculated from the data shown in Figure 4.

We observed a large spread in the measured bleedthrough coefficients in the data from live cells and beads. The bacterial data showed less variability, probably because each measurement was obtained from an entire field of view, making it less susceptible to imaging artefacts and providing a measure of the average of many more molecules than those found in a single cell or bead. This spread was especially clear in the fluorophores and channels with the highest bleedthrough. Since we will unmix the signals obtained in single cells, we decided to use the average coefficients calculated from the live cell measurements.

### Shading

After visually inspecting the data, we observed that the images were affected by shading. To quantify shading, we prepared fluorescent protein solutions and imaged them. As shown in Figure 5, the shading was substantial. Fluorescence intensity counts ranged from 65% to 125% of the average, which resulted in up to a 2-fold change between pixels within a single field of view. The counts tended to increase from the bottom left to the top right of each quadrant, although the exact distribution varied between channels.

**Figure 5.**
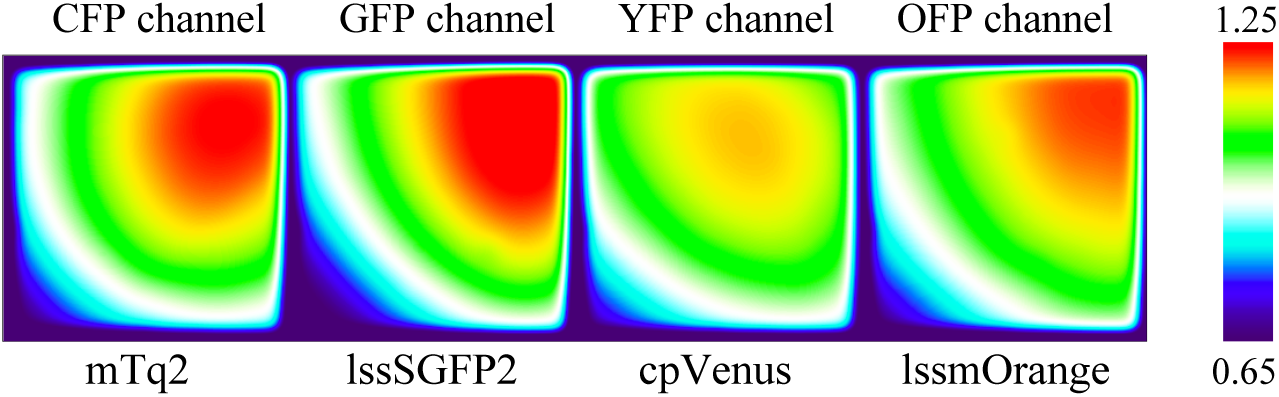
Shading pattern in the 4 channels. Solutions from each fluorescent protein were prepared and imaged in the multi-channel system. Fluorescence intensity counts were normalized to the average fluorescence intensity of the field of view. For each channel, a representative image of the fluorescent protein with highest spectral overlap is shown.

The observed differences in the shading distribution between channels would alter the fluorescence intensity ratio calculated from a pixel in any two given channels. Therefore, we decided to reanalyze the live cell data used for the bleedthrough calculation after applying the shading correction. By pairing the bleedthrough percentages per cell before and after the correction, we observed changes in individual measurements of up to 10%, as shown in Figure 6. However, these individual differences cancelled each other and the average bleedthrough percentages were not affected, as shown in Table 3.

**Figure 6.**
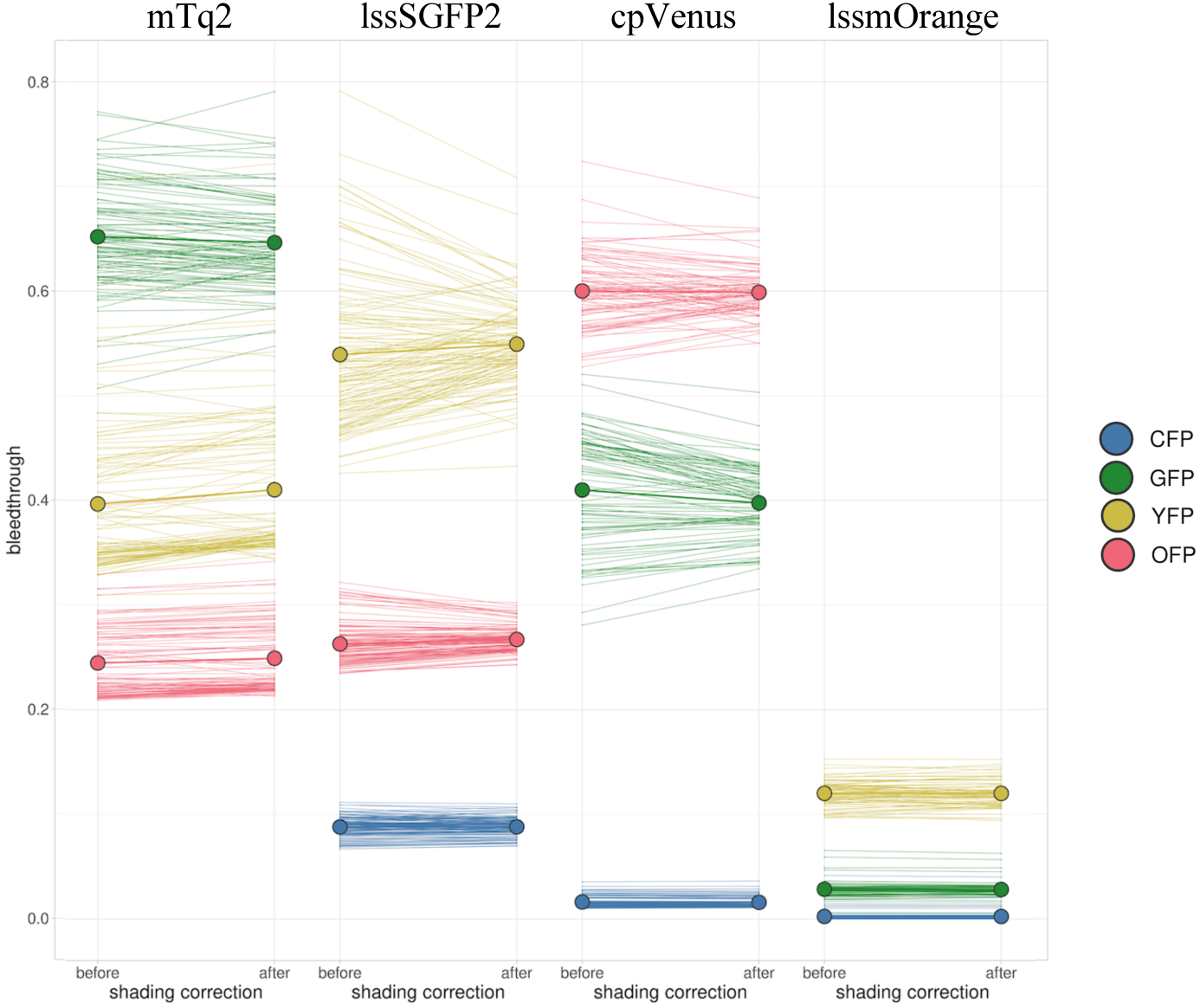
Effect of shading correction in the calculated bleedthrough per channel in live cell measurements. The bleedthrough for each FP in every channel was recalculated after correcting the images for shading. Each thin line connects a single cell measurement before and after the correction, and the thick lines connect the average of all single cell measurements, indicated by a thick dot. Data was plotted using SuperPlotsofData [25].

**Table 3.** Effect of shading correction in the calculated bleedthrough per channel in live cell measurements. The average and standard deviation of the measurements per FP per channel were calculated from the data shown in Figure 6.

| shading correction |  |  |  |  |  |  |  |
| --- | --- | --- | --- | --- | --- | --- | --- |
| before |  |  |  | after |  |  |  |
| mTq2 |  |  |  | mTq2 |  |  |  |
| channel | n | mean | SD | channel | n | mean | SD |
| CFP | 101 | 1 | 0 | CFP | 101 | 1 | 0 |
| GFP | 101 | 0.65 | 0.05 | GFP | 101 | 0.65 | 0.04 |
| YFP | 101 | 0.4 | 0.07 | YFP | 101 | 0.41 | 0.07 |
| OFP | 101 | 0.24 | 0.03 | OFP | 101 | 0.25 | 0.04 |
| lssSGFP2 |  |  |  | lssSGFP2 |  |  |  |
| CFP | 125 | 0.09 | 0.01 | CFP | 125 | 0.09 | 0.01 |
| GFP | 125 | 1 | 0 | GFP | 125 | 1 | 0 |
| YFP | 125 | 0.54 | 0.07 | YFP | 125 | 0.55 | 0.04 |
| OFP | 125 | 0.26 | 0.02 | OFP | 125 | 0.27 | 0.01 |
| cpVenus |  |  |  | cpVenus |  |  |  |
| CFP | 87 | 0.02 | 0.01 | CFP | 87 | 0.02 | 0.01 |
| GFP | 87 | 0.41 | 0.05 | GFP | 87 | 0.4 | 0.03 |
| YFP | 87 | 1 | 0 | YFP | 87 | 1 | 0 |
| OFP | 87 | 0.6 | 0.03 | OFP | 87 | 0.6 | 0.02 |
| lssmOrange |  |  |  | lssmOrange |  |  |  |
| CFP | 73 | 0 | 0.01 | CFP | 73 | 0 | 0.01 |
| GFP | 73 | 0.03 | 0.01 | GFP | 73 | 0.03 | 0.01 |
| YFP | 73 | 0.12 | 0.01 | YFP | 73 | 0.12 | 0.01 |
| OFP | 73 | 1 | 0 | OFP | 73 | 1 | 0 |

### Generation of stable cell lines

Following a similar approach described in a previous study [18] and given the added complexity of the imaging system, we decided to generate stable HEK-CaSR cell lines using the PiggyBac transposon system [20]. Since the two reporters have different designs and read-outs, we decided to have the Gαi1 FRET sensor in one plasmid, and the ERK-KTR with the nuclear marker in another plasmid, in order to control the relative expression of both constructs. We generated stable cell lines that expressed both constructs and used fluorescence-activated cell sorting (FACS) to isolate 4 populations with different expression levels of each construct (data not shown). For the FRET pair, we set two gates based for “medium” and “high” counts of cpVenus, and for the H2A-KTR pair we used lssSGFP2 intensity to set gates for “low” and “medium” intensity. From these populations, and upon inspection of the fluorescence intensities per channel, we decided to continue with the one with higher expression of both constructs and performed serial dilutions to obtain monoclonal populations. From the expanded cell populations, we found one cell line that expressed both constructs and showed homogeneous expression among the single cells within the population, as can be observed in Figure 7A.

**Figure 7.**
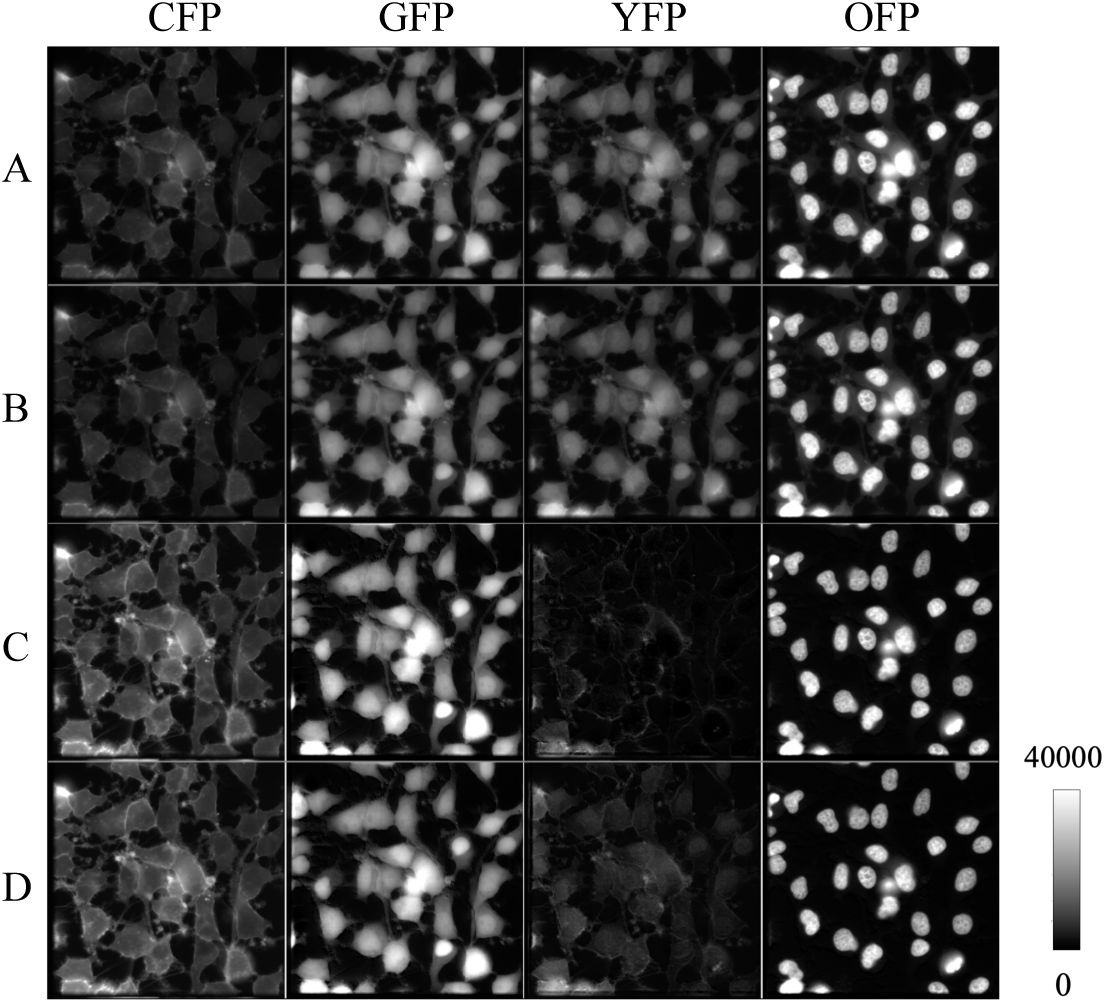
Effect of shading and unmixing correction over the fluorescent signals in the four imaged channels. Stable cell line expressing Gi1 FRET sensor, ERK-KTR lssSGFP2, and H2A-lssmOrange was imaged, and the images were corrected for shading and for bleedthrough using two different sets of coefficients. A: Raw images. B: Shading-corrected images. C-D: Shading-corrected images corrected for bleedthrough with unmodified and adjusted coefficients.

### Corrected unmixing

After unmixing the shading-corrected images, shown in Figure 7B-C, it became clear that the nuclear intensity counts of the unmixed lssSGFP2 and lssmOrange were considerably higher than those of mTq2 and cpVenus. This is partially explained by the relative localization of the constructs, with the FRET sensor localizing predominantly in the membrane. However, we found negative pixels in the unmixed cpVenus image, which resulted from the combination of an “imperfect” unmixing matrix, the relatively low YFP counts, and the bleedthrough from the other, brighter, signals. To address this, we decided to reduce the coefficients in the unmixing matrix for the YFP channel. For this, we went back to the single cell measurements in the YFP channel and we decided to calculate the average of the lowest 50%, 10% and 10% bleedthrough values of mTq2, lssSGFP2 and lssmOrange respectively. This resulted in coefficients about 12-20% lower than the previously calculated, as shown in Table 4. After applying the modified matrix, the unmixed cpVenus image does not show negative values, as shown in Figure 7D, but we observe residual nuclear counts in the YFP channel from an insufficient subtraction of the lssSGFP2 and lssmOrange.

**Table 4.**
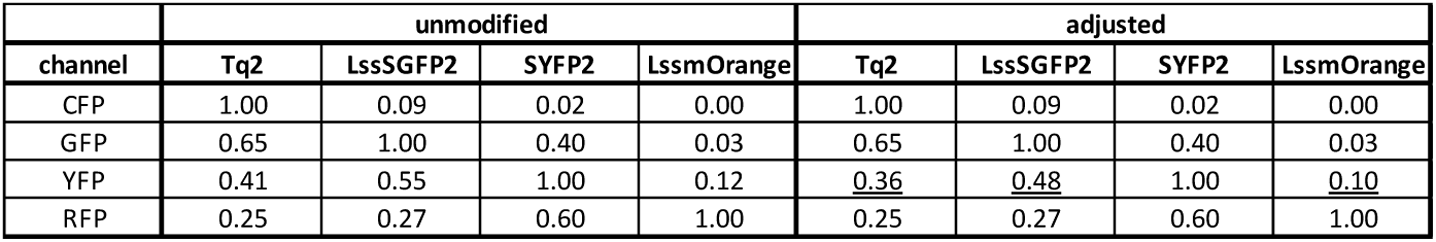
Comparison between unmodified and adjusted bleed-through coefficients. The newly calculated average bleed-through coefficients of mTq2, lssSGFP2 and lssmOrange in the YFP channel are shown underlined.

### Background subtraction

To remove the background fluorescence in the images, we first tried the “Subtract background” tool from FIJI. It is commonly used to correct for uneven background by applying the “rolling ball” algorithm with a radius equivalent to the largest foreground object (i.e. the nucleus). We found very large disparity in the calculated background values within a single field of view, as exemplified in Figure 8B with subtracted values ranging between 100 and 900 counts. This is a direct effect of cell density, as confirmed in Figure 8E where a group of cells are very closely together and the algorithm considers the background to be the area with lower counts between the nuclei. In both cases the real background is around 1000 counts.

**Figure 8.**
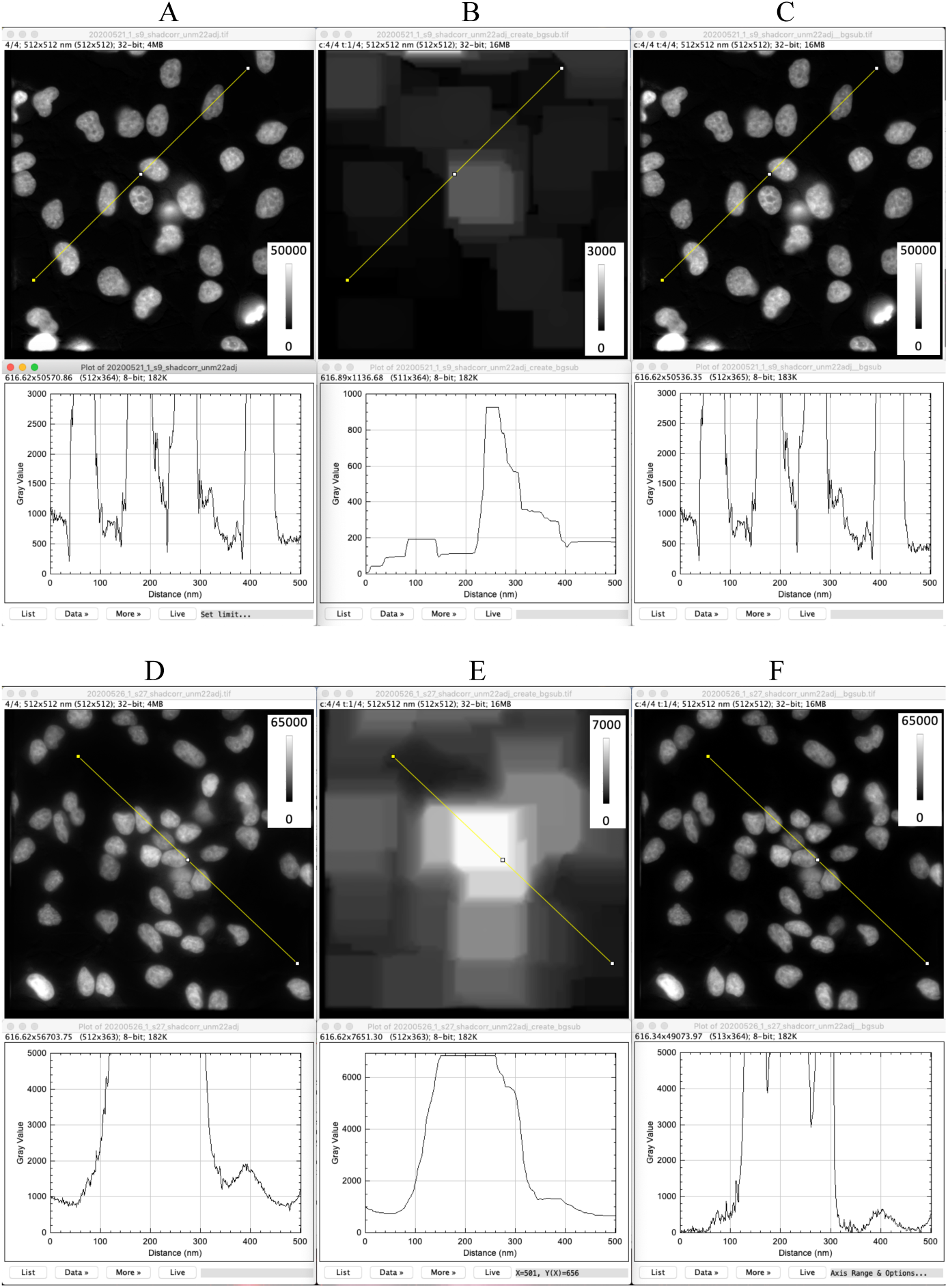
Effect of cell density over rolling ball background subtraction. The effect of background subtraction with a rolling ball radius of 55 pixels nuclear channel of two different fields of view with low and high cell density are shown. Per field of view, the top row shows images while the bottom row shows the counts profile of the shown ROI. A-C: Field of view with low cell density. D-F: Field of view with high cell density. A, D: Raw images. B, E: Background to be subtracted. C, F: Images after background subtraction.

Given the limited control we have over cell distribution in a field of view, we decided to not use this tool as it would directly modify the images intensity values in an uncontrolled manner. Instead, we decided to manually draw a background ROI and subtract this value from the entire field of view. This approach does not correct for uneven background fluorescence, but that seemed unnecessary considering the low variability within the different fields of view.

### Segmentation and tracking

To quantify the ERK and Gi activities, we used StarDist [26] and a custom-made CellProfiler [27] pipeline for segmentation, measurement of intensity and shape features, and tracking. First, we segmented the nuclei using StarDist, then, in CellProfiler, we used these nuclei masks as seeds to obtain cellular masks. A representative example is shown in Figure 9A. Later, using CellProfiler, both objects were tracked over the timelapse based on overlap between consecutive images. Figure 9B shows tracked nuclei at 3 different timepoints.

**Figure 9.**
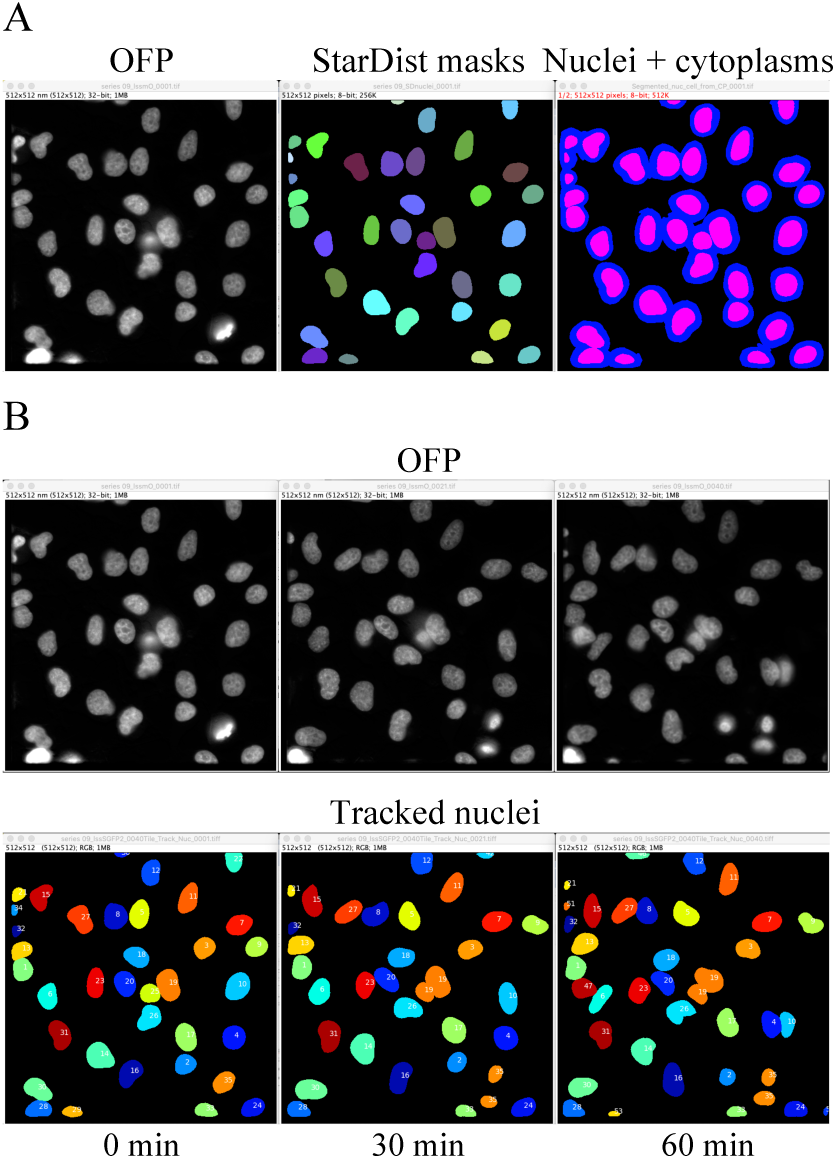
A: Segmentation of nuclei and cytoplasms. Left, raw lssmOrange image. Center, segmented nuclear mask by StarDist. Right, in magenta, nuclear mask, in blue, cytoplasmic ROI. B: Tracking of nuclei over the timelapse. Top, raw lssmOrange image. Below, tracked nuclei.

### Seeding conditions

FBS contains multiple growth factors that are essential for cell growth and division, such as EGF, IGF, PDGF, TGF, and FGF2. Many of these growth factors activate ERK signaling causing transient, sustained, and oscillatory responses in a concentration dependent manner [28–30]. In order to reduce basal ERK activity, we decided to serum-starve the cells for at least 2 hours. In addition, we needed to starve the cells from calcium for 20 min, as it is the primary endogenous ligand for CaSR.

We started by seeding the cells with full growth medium the night before imaging. HEK-CaSR cells form “clumps” and their nuclei are very large relative to the cytoplasm, which complicates the segmentation and tracking. To address this, we mixed the cells with untransfected HEK-CaSR cells, which resulted in a partial improvement, as shown in Figure 10. As previously reported in another study [18], we needed to reduce the basal ERK and CaSR activities in the cells to have an assay window to observe changes in ERK and G protein following CaSR activation. Replacing the full growth medium with 0.5 mM Ca^2+^ microscopy medium, to starve the cells from serum and calcium simultaneously, resulted in immediate translocation of the ERK-KTR to the cytoplasm suggesting a strong stress response, as shown in Figure 11. Only after about 1 h the KTR relocalized to the nucleus.

**Figure 10.**
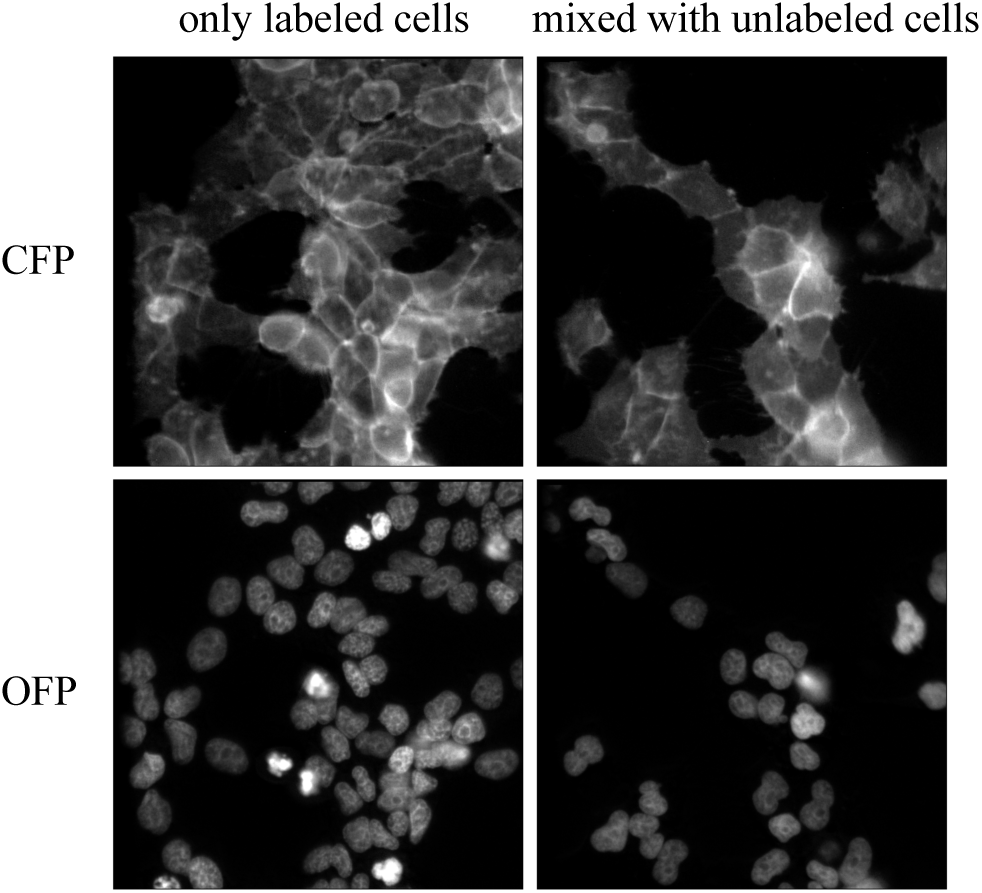
Effect of mixing generated stably expressing cells with untransfected HEK-CaSR cells. Representative images of the CFP and OFP channels, showing the stably expressing cells, seeded alone or in combination with untransfected HEK-CaSR cells at a 1:1 ratio.

**Figure 11.**
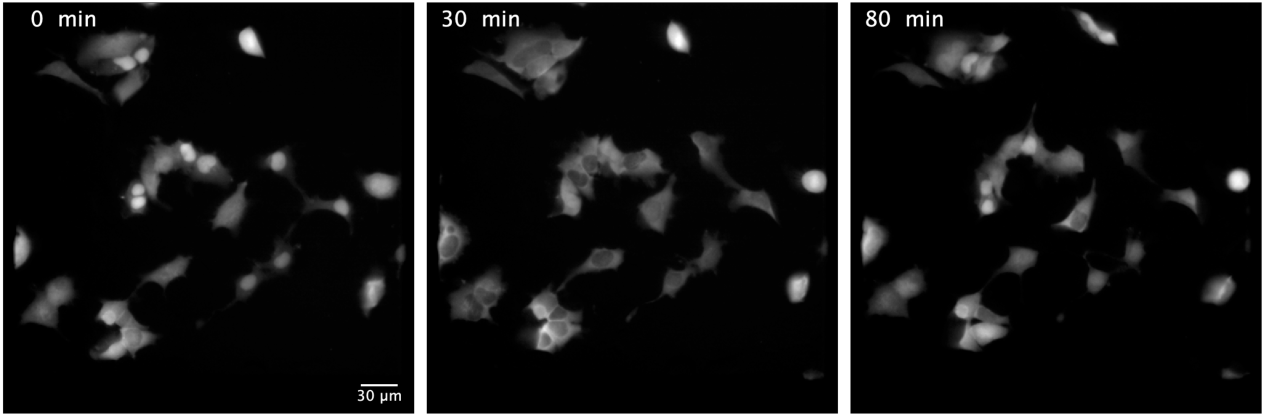
Effect of replacing the medium over localization of ERK-KTR. Representative images of ERK-KTR localization in the cells over time after medium replacement.

To avoid replacing the medium prior to imaging and to minimize the clumpiness of the cells, we seeded the cells 2 h before imaging with serum-free 0.5 mM Ca^2+^ DMEM. However, the cells did not attach, and increasing the Ca^2+^ concentration to the physiological level of 1.8 mM improved attachment in only a small fraction of the cells (data not shown). We found that HEK-CaSR cells fully attach after 5 h under typical culture conditions, in full growth medium, as shown in Figure 12. Subsequently, we tested seeding the cells with serum-free media, including FluoroBrite (FB), 1.8 mM Ca^2+^ Microscopy Medium (MM), and a 1:1 combination of FB and 0 mM Ca^2+^ MM (0.9 mM Ca^2+^ FB/MM). FB is specially formulated for imaging due to lower autofluorescence and can be used for routine cell culture when supplemented with FBS, while MM is formulated to maintain cell homeostasis for short periods of time during imaging. However, 5h after seeding, the cells are rounder compared to the attached cells with the full growth medium., as shown in Figure 12, especially the low-calcium FB/MM mix. Comparison of these seeding conditions also showed clear differences in background fluorescence, with MM showing the lowest counts in all channels, followed by the 1:1 mix with FB, as can be seen in Figure 13. Considering these results, we decided to seed the cells the night before in full growth medium, and to replace the medium 2h prior to imaging with low-calcium MM or FB/MM. Even though MM provided the lowest background, we observed that some of the cells detached very easily upon pipetting (data not shown). Therefore, we decided to use the FB/MM mix for serum and calcium starvation prior to imaging.

**Figure 12.**
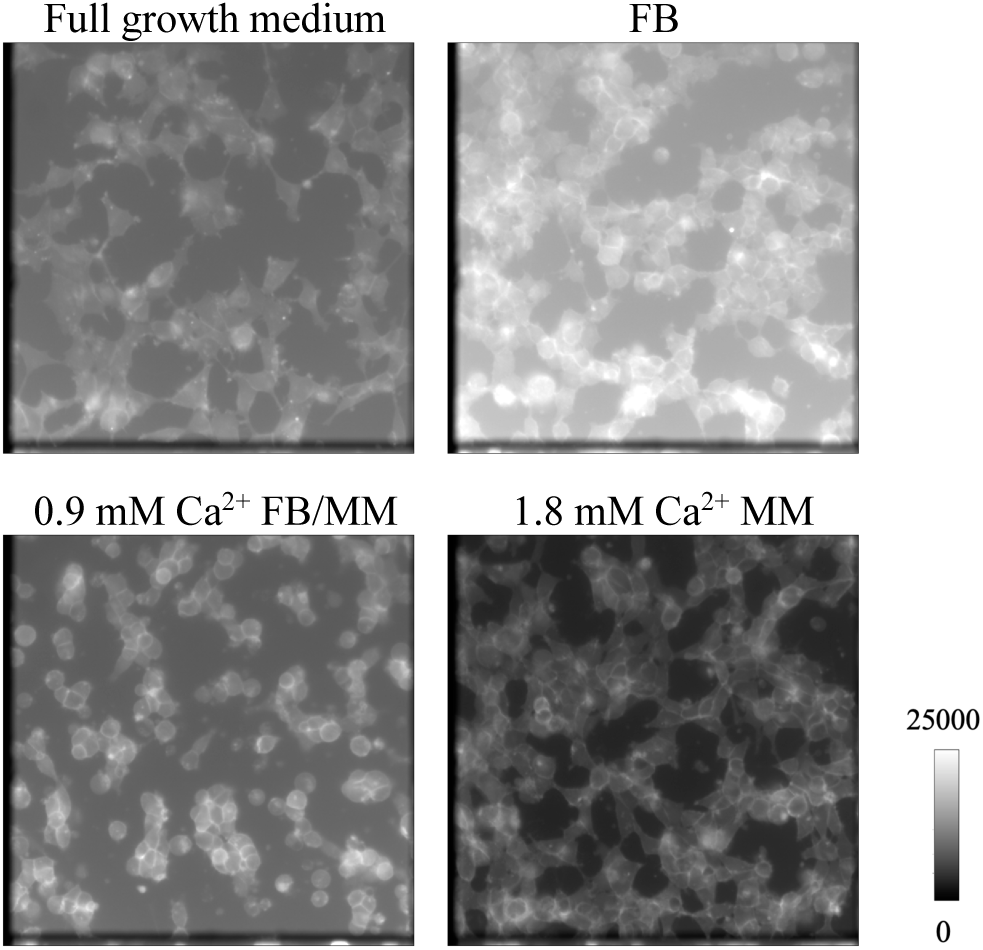
Effect of using different media during seeding HEK-CaSR cells prior to imaging. Representative images of stably expressing HEK-CaSR cells 5h after seeding with the following media: Full medium (DMEM + 10% FBS), FluoroBrite (FB), 1.8 mM Ca^2+^ MM, and 0.9 mM Ca^2+^ FB/MM.

**Figure 13.**
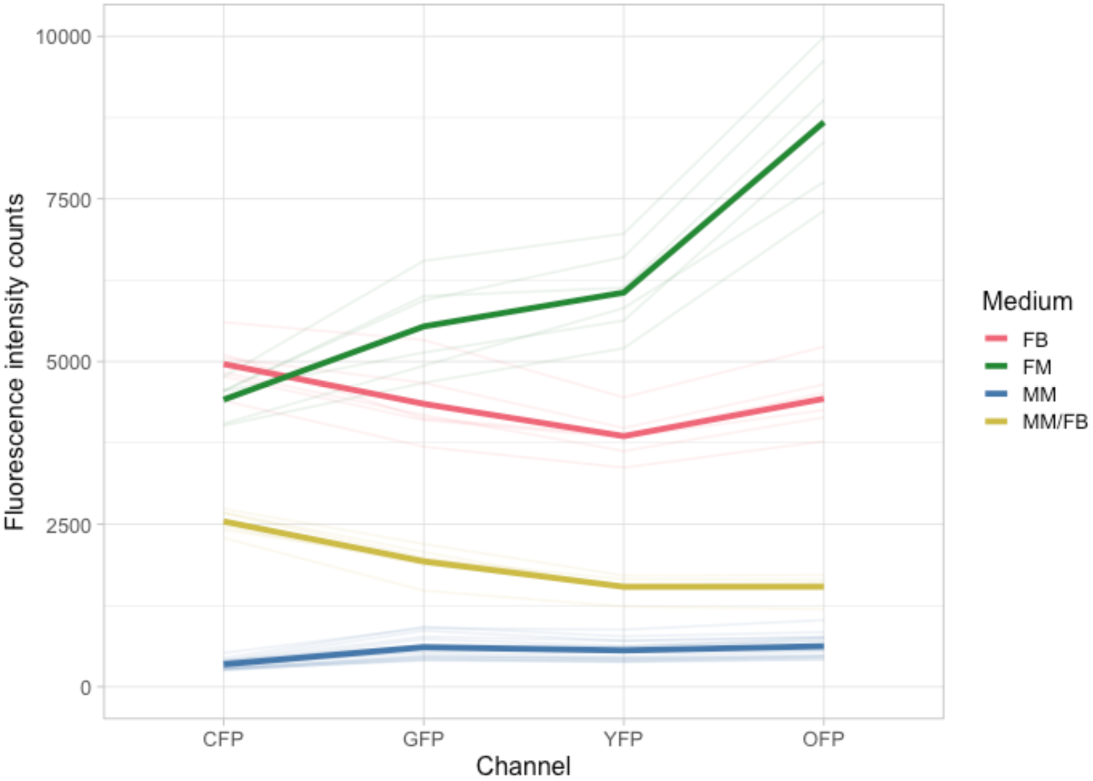
Comparison of the fluorescence of different culture media. Thick colored lines show the mean and thin colored lines show the standard deviation of the fluorescence intensity counts in the 4 channels for each of the tested media.

### Effect of calcium stimulation

To test the effect of stimulating the CaSR with Ca^2+^ on the ERK and Gi activities, we treated the cells with either 10 mM Ca^2+^ or medium (0.9 mM Ca^2+^ FB/MM) as control. As can be observed in Figure 14A, the average ERK KTR C/N ratio started to increase rapidly after addition of calcium and it continued to increase for up to 50 minutes when the experiment was stopped. The medium control showed a lower and slower increase that stabilized about 25 min after treatment.

**Figure 14.**
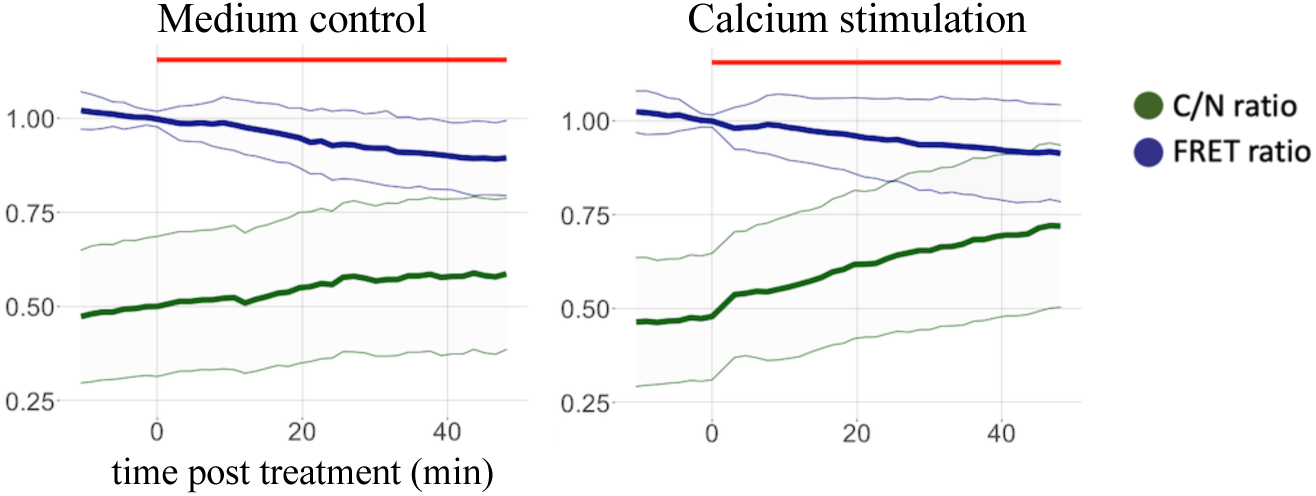
Effect of calcium stimulation over ERK and Gi activity measurements. Cells were treated with medium (medium control) or 10 mM Ca^2+^ (calcium stimulation) for the time indicated with the red line. Gi FRET ratios, indicated in dark green, are normalized by dividing by the value prior to treatment. ERK KTR C/N ratios are indicated in dark blue. Dots and thin lines show the mean and standard deviation respectively. The number of cells was 157 for medium control and 234 for calcium stimulation.

In terms of Gi activity, measured by the ratio between the cellular counts in the YFP and CFP channels, there was no clear difference between the average ratios in the two tested conditions directly upon stimulation. G protein activation is reflected by an immediate FRET ratio decrease that is sustained over several minutes [10]. Therefore, G protein activation would be reflected by a clear decrease at the first measured timepoint after stimulation, which would remain constant for several minutes. Instead, we found a slow constant decrease in the YFP/CFP ratio over time, both for calcium-treated and medium-treated conditions. The observed average YFP/CFP ratio kinetics appears to share a similar trend as that of the KTR, although inverted. This could be explained by a decrease in the cellular YFP counts as a result from residual bleedthrough from lssmOrange and/or lssSGFP2, both of which decrease overtime as shown in Figure 15A. Upon further exploration of the average fluorescence counts in the nuclear masks, shown in Figure 15B, we found similar dynamics for the average lssmOrange and lssSGFP2 counts overtime. Since the nuclear marker is relatively stable, this is probably the effect of residual bleedthrough from the dynamic lssSGFP2. The cytoplasmic counts, shown in Figure 15C, show similar dynamics for GFP, CFP and YFP.

**Figure 15.**
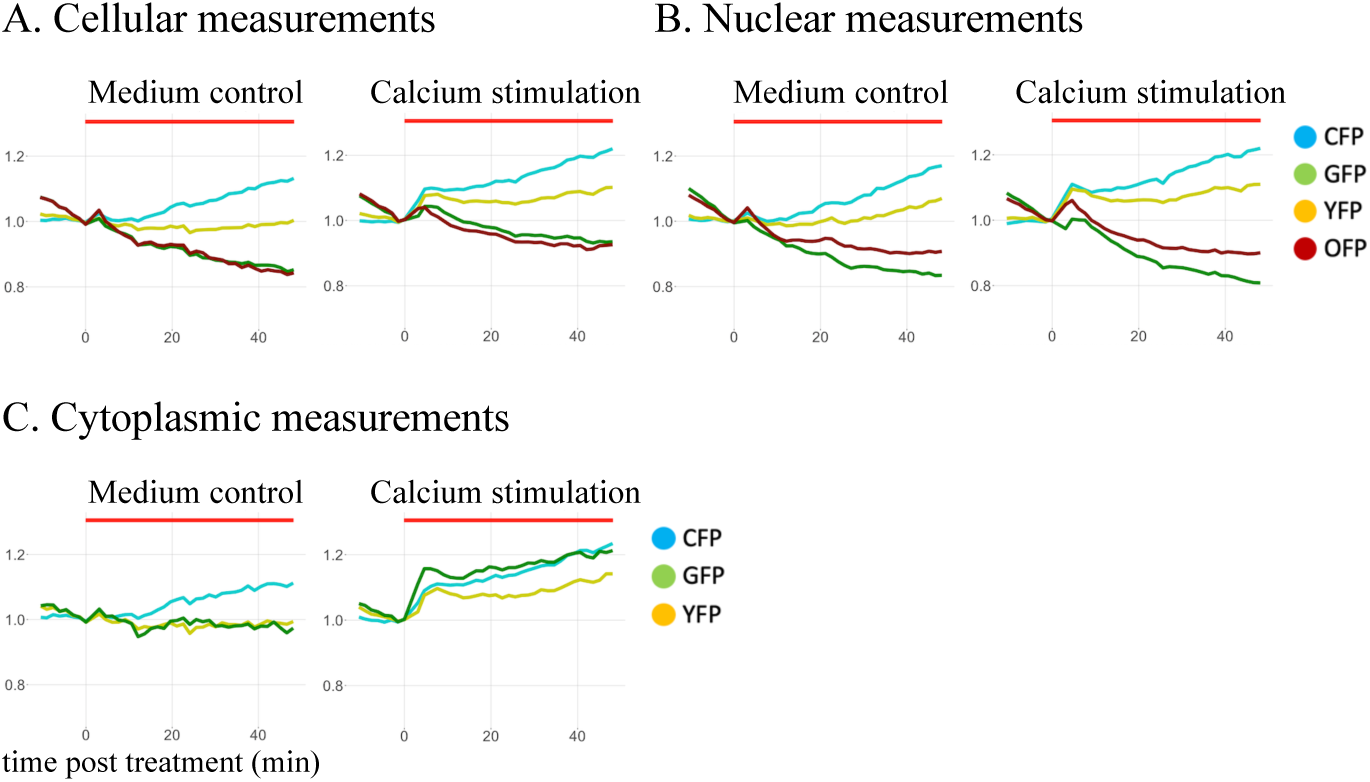
Average intensity measurements per channel per compartment. Intensity counts from all 4 channels were measured in the cellular (A), nuclear (B), and cytoplasmic (C) compartments in the cells from Figure 14, and normalized by dividing by the value prior to treatment. Counts in the CFP, GFP, YFP, and OFP channels are shown in light blue, green, yellow, and red respectively. Dots indicate the mean.

When single cell measurements are examined, we observe similar trends between nuclear GFP and OFP, and between cytoplasmic GFP, CFP and YFP, as shown in Figures 16. Unsurprisingly, the measured FRET ratios in these single cells do not show stable results, and no clear consistent difference is observed between the nuclear, cytoplasmic, and cellular FRET ratios, as shown in Figure 17.

**Figure 16.**
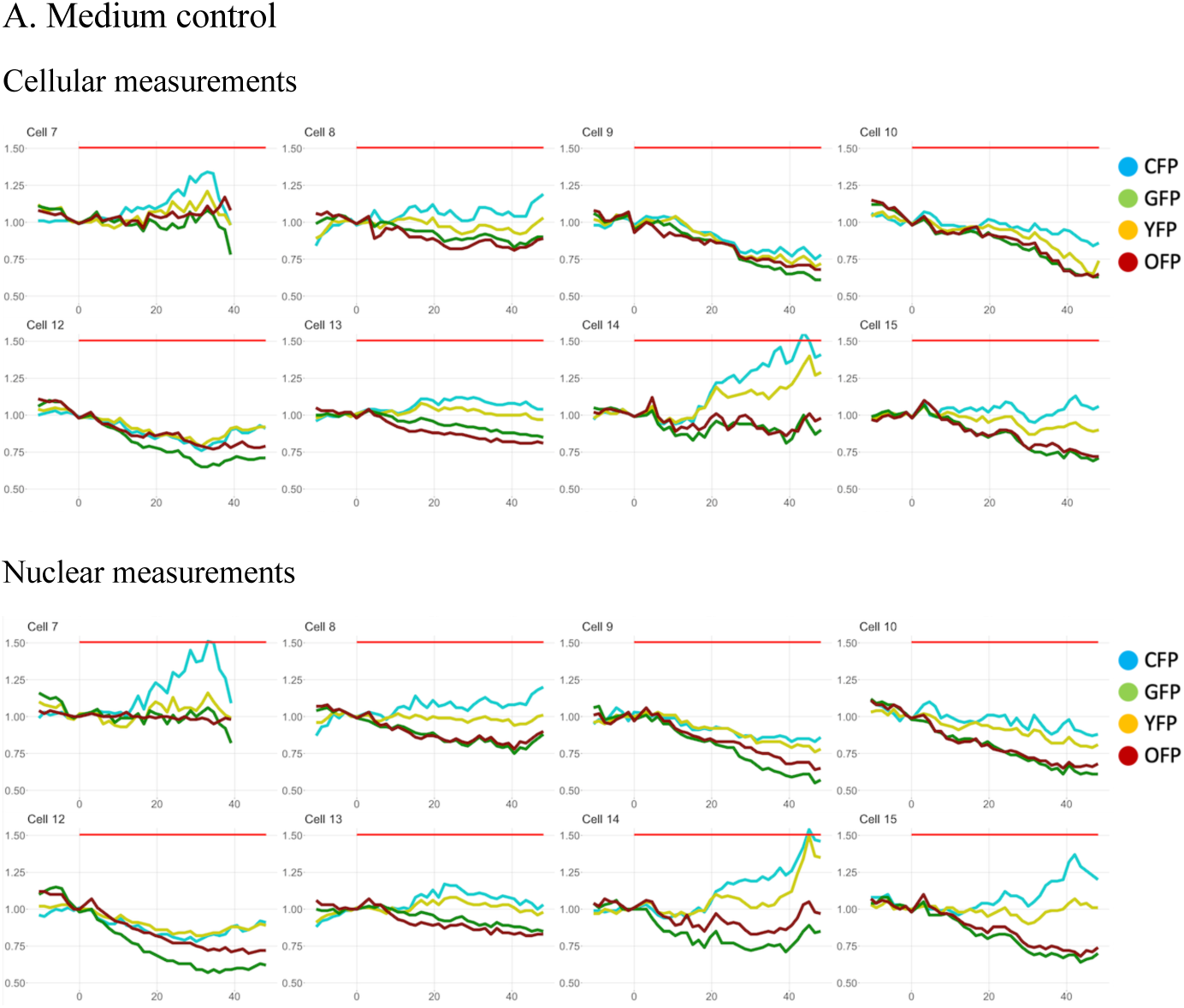
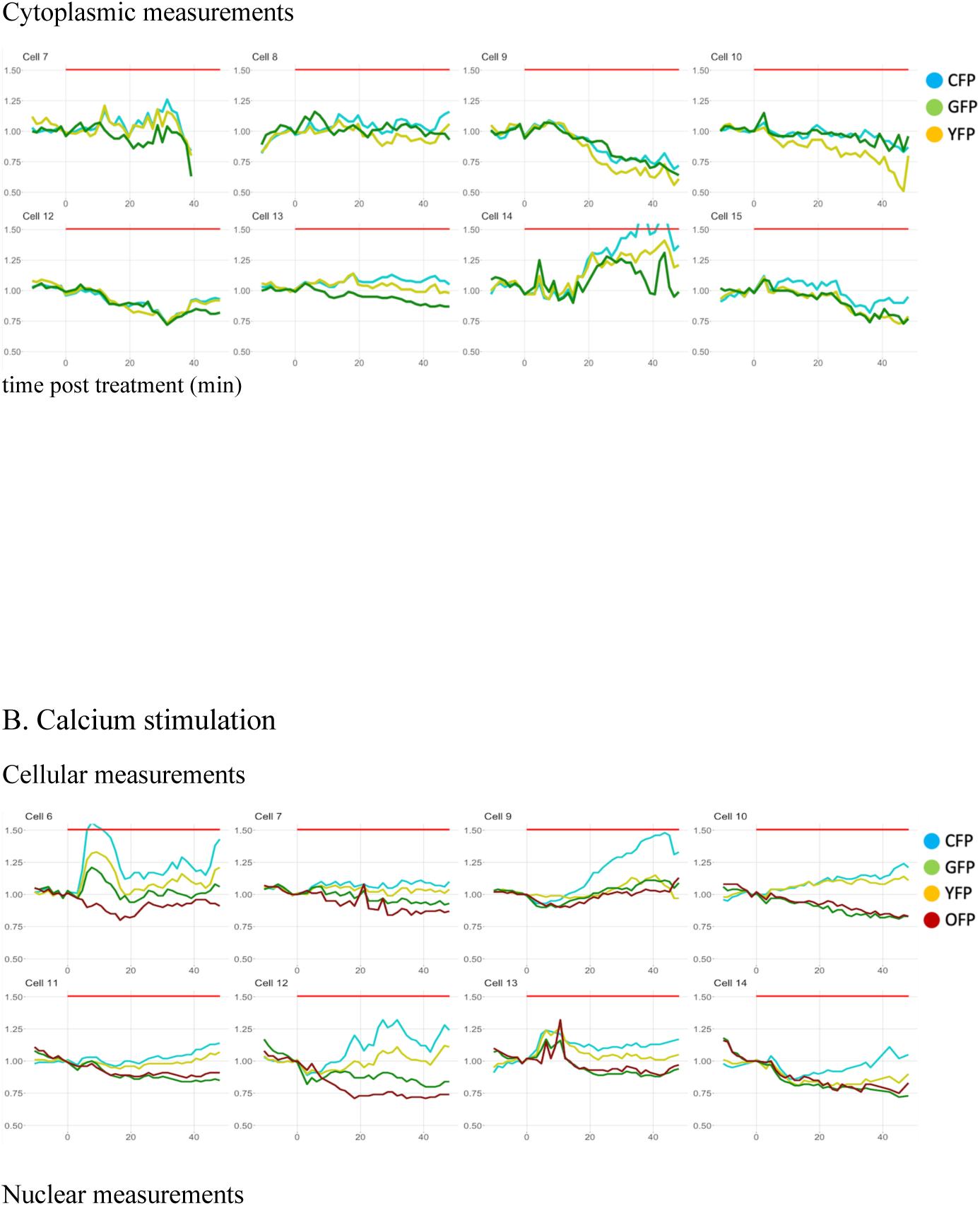
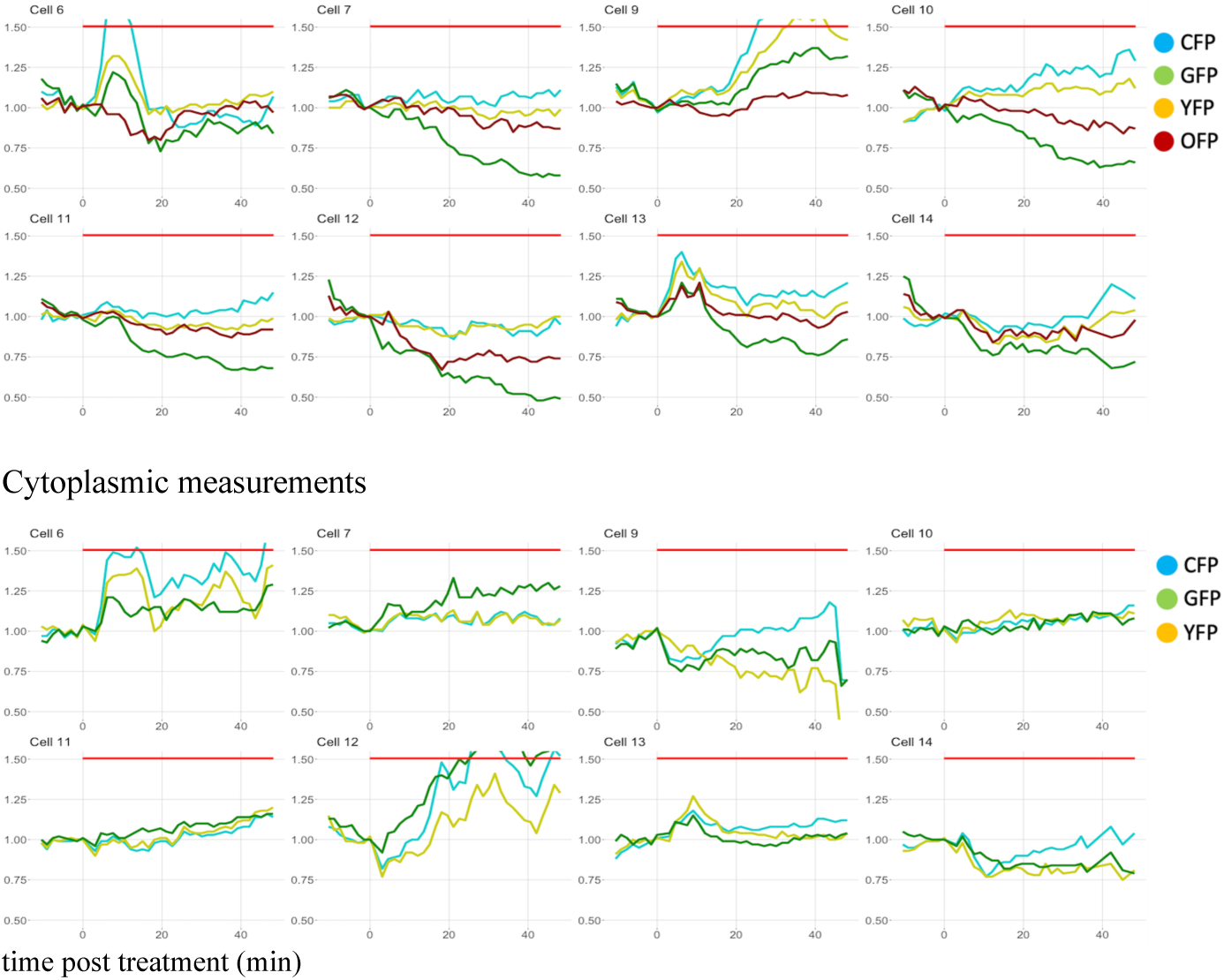
Single cell intensity measurements per channel per compartment. Individual intensity counts from all 4 channels in the cellular, nuclear, and cytoplasmic compartments are shown for a subset of cells from Figure 14. Counts in the CFP, GFP, YFP, and OFP channels are shown in light blue, green, yellow, and red respectively.

**Figure 17.**
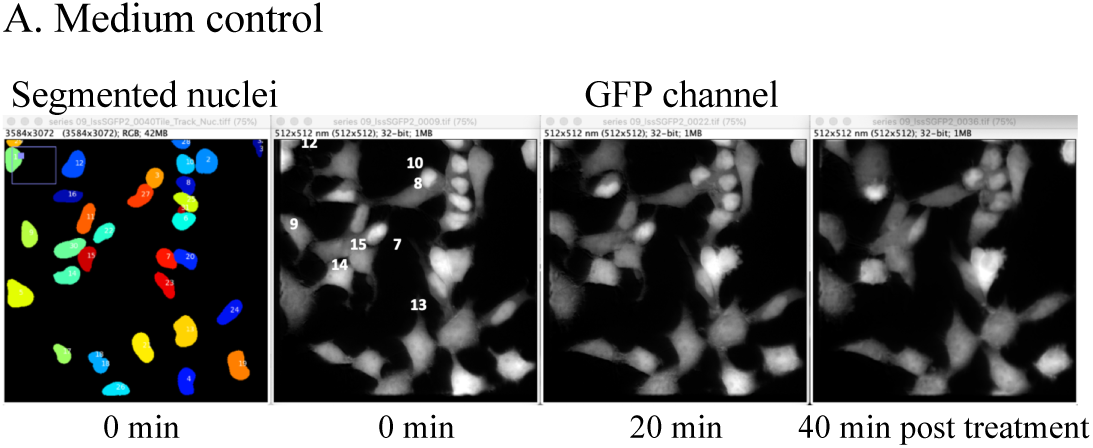
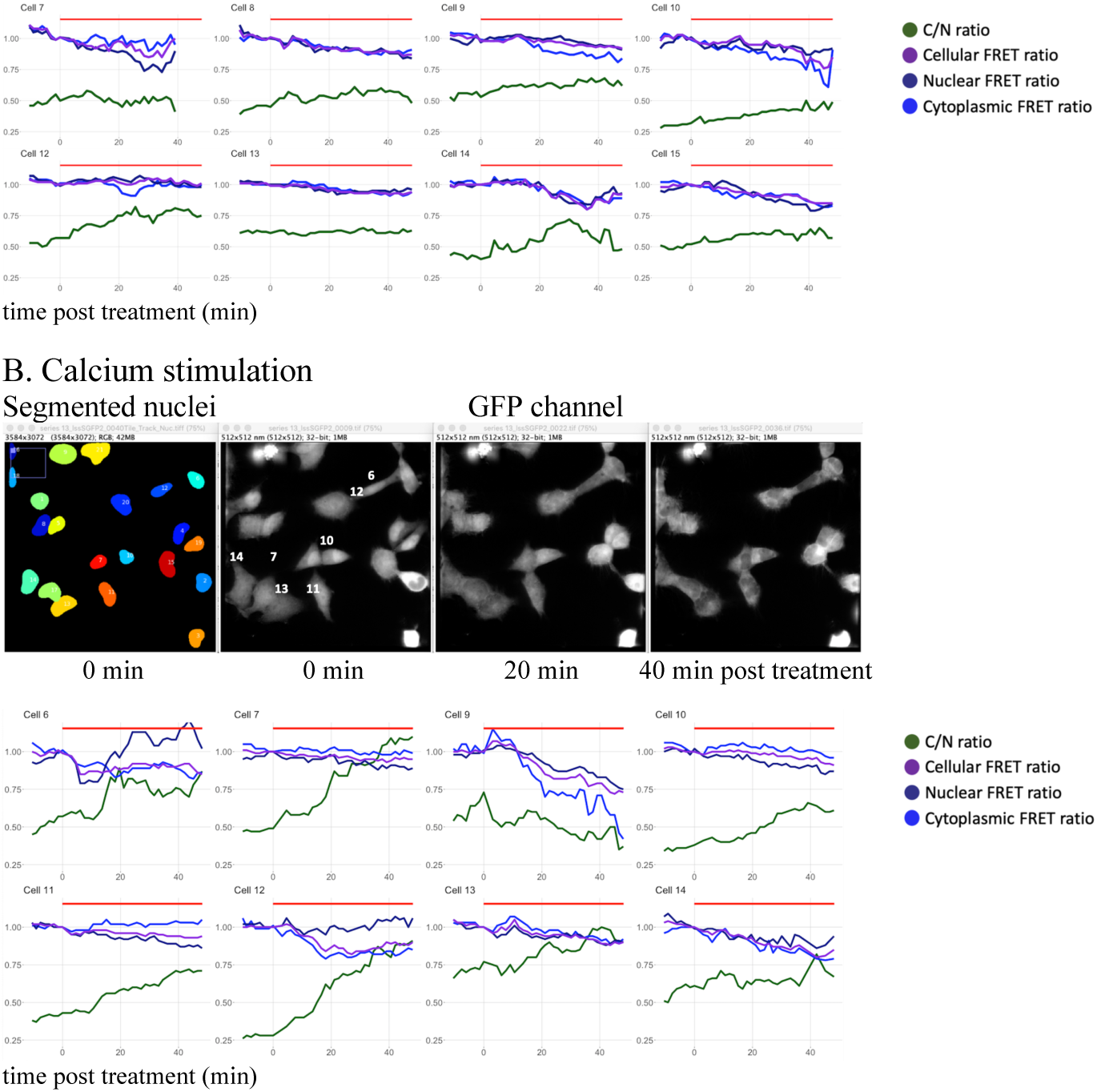
Single cell ERK and FRET measurements. Cells treated with medium control (A) or stimulated with calcium (B) were tracked over time and the C/N and ERK ratios were calculated. Top panel shows from left to right: Segmented nuclei at time point 0 and GFP channel images at time points 0, 20, and 40 min post treatment. Bottom panel shows the C/N ratios (in dark green) and FRET measurements in the different compartments (cytoplasmic in blue, cellular in purple, and nuclear in dark blue) for the single cells indicated in the top panel. FRET ratios are normalized by dividing by the value prior to treatment.

These observations altogether suggest that the unmixing of the signals was not sufficient or precise enough, which makes very challenging to obtain a clear FRET ratio change in the current setup, considering the large differences in intensity between the FRET pair and the lssSGFP2/lssmOrange, the high bleedthrough from LssSGFP2, the dynamicity of the LssSGFP2-tagged KTR probe, and the relatively low dynamic range of the FRET pair.

On the other hand, we observed clear translocation and the corresponding change in the quantified C/N ratio in those same cells upon calcium stimulation. In medium control treated cells, some cells showed a modest translocation compared to the calcium stimulated ones, as was already indicated by the average shown in Figure 14. For these reasons, we decide to focus solely on the ERK-KTR measurements.

Next, we wanted to validate these results by using various concentrations of calcium and in the presence of CaSR inhibitor NPS-2143. In the absence of NPS-2143, addition of medium and of 3 mM, 5 mM, and 10 mM Ca^2+^ resulted in increased C/N ratio overtime, as shown in Figure 18. On the other hand, when NPS-2143 was added before stimulation, stimulation with 10 mM Ca^2+^ showed a much clearer increase than the other conditions. As we noticed that the average C/N ratios prior to stimulation were different between the tested conditions, we normalized the data to compare the different trajectories over time. In conditions without NPS-2143, we observed now that the cells stimulated with 5 and 10 mM Ca^2+^ showed higher C/N ratios up to an hour after stimulation. In contrast, upon NPS-2143 pre-treatment, the cells treated with 10 mM Ca^2+^ showed consistently higher increased C/N ratios up to 90 min after stimulation.

**Figure 18.**
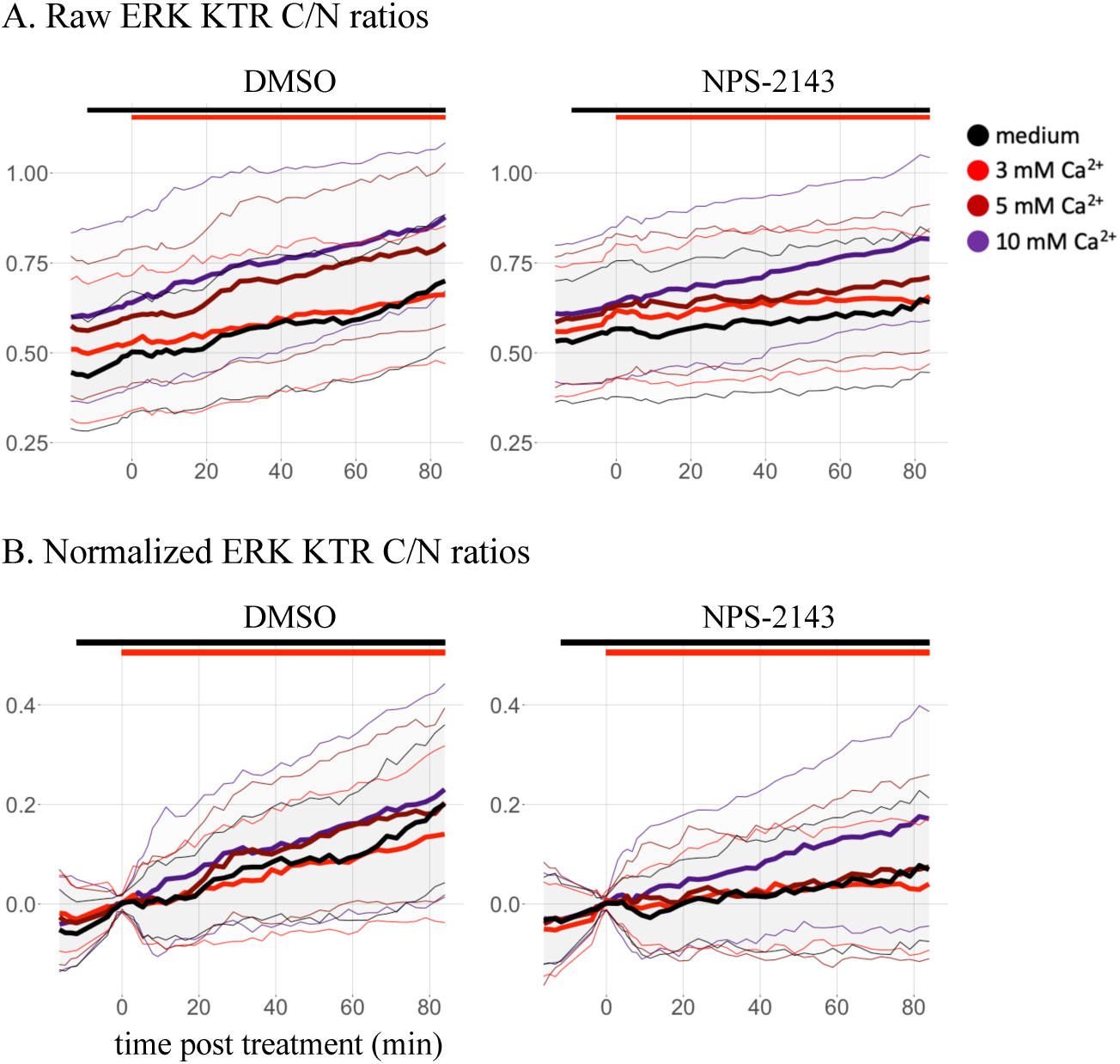
Effect of stimulation with multiple calcium concentrations on ERK and dependency to CaSR. A: Raw data. B: Normalized data to C/N ratio prior to treatment. Cells were pre-treated with vehicle (DMSO) or CaSR inhibitor (1 μM NPS-2143), and stimulated with medium (in black), 3 mM Ca^2+^ (in red), 5 mM Ca^2+^ (in dark red), or 10 mM Ca^2+^ (in purple). Black and red bars indicate pre-treatment and stimulation respectively. Dots indicate the mean and thin lines show the standard deviation. The numbers of cells per trajectory ranged between 71 and 138 cells.

Given the observed differences in start C/N ratios between the tested conditions, we decided to determine whether this variability influenced the measurements. Therefore, we looked at the distribution of the C/N ratios before stimulation and we found that they are spread between 0.20 and 1.10, as shown in Figure 19. We then split the data into three groups: low (up to 25th percentile), medium (between 25th and 75th percentile), and high (above 75th percentile). The 3 grouped trajectories are shown in Figure 20, revealing that for any given condition the amplitude of the measured response per subpopulation decreases when the start ratio increases. In addition, we observed a possible positive correlation between the calcium concentration and the maximum C/N ratio, as well as a possible inhibitory effect of NPS-2143. Upon comparison of the low and medium subpopulations in vehicle pre-treated cells, which are the most responsive, we observed the largest decreased response in the cells stimulated with 10 mM Ca^2+^. Interestingly, in NPS pre-treated cells, a similar phenomenon is observed but for the 5 mM Ca^2+^.

**Figure 19.**
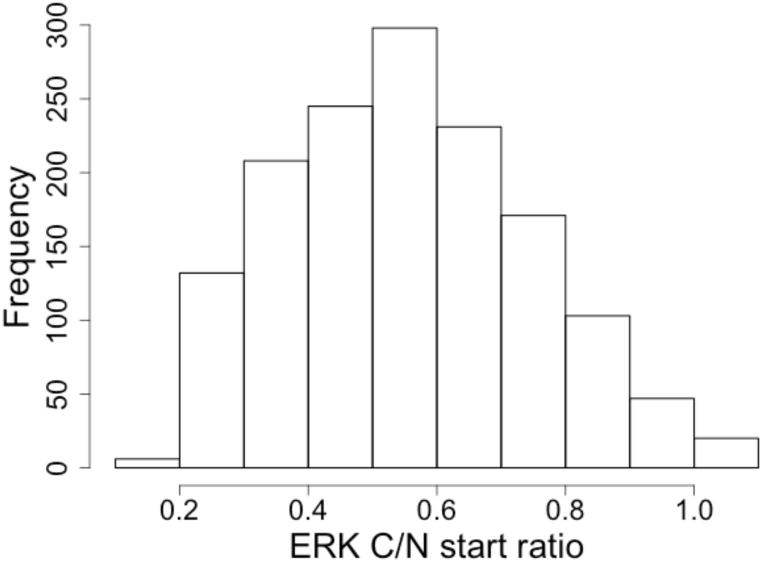
Distribution of start ERK C/N ratios. Frequency of the average C/N ratios of ERK before treatment from all single cells tracked in the experiments.

**Figure 20.**
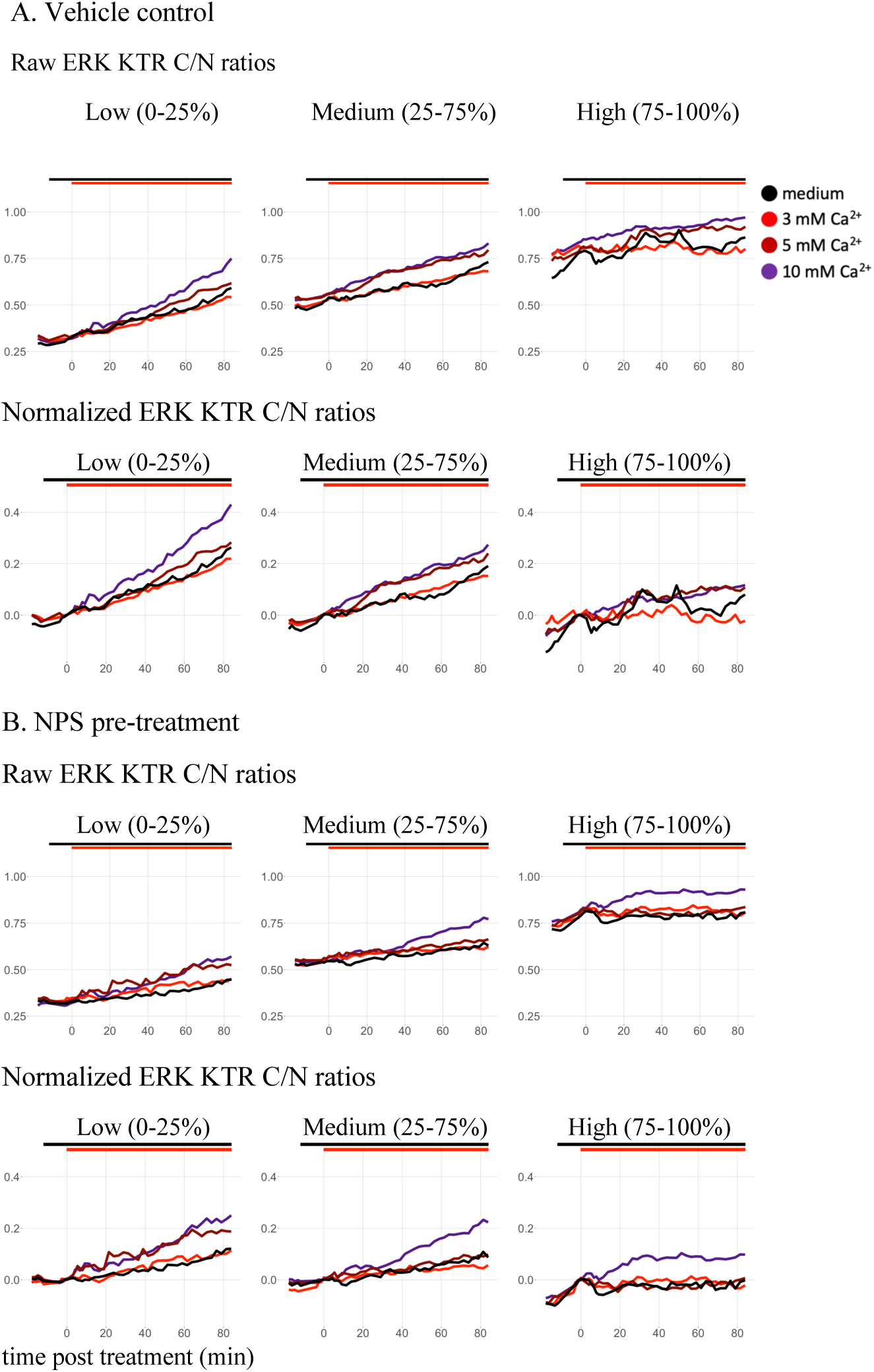
Influence of start activity over ERK responses over time upon calcium stimulation. A: Vehicle control. B: CaSR inhibitor. Raw (top) and normalized to C/N ratio prior to treatment (bottom). Data from Figure 18 was divided in 3 groups, based on the start ratio per cell. Cells were pre-treated with vehicle DMSO or with 1 μM NPS-2143, and stimulated with medium or 3, 5, or 10 mM Ca^2+^. Black and red bars indicate addition of pre-treatment and stimulation respectively. Black, red, dark red, and purple colored trajectories correspond to treatments with medium, 3, 5, and 10 mM Ca^2+^ respectively. Dots indicate the mean and thin lines show the standard deviation. The numbers of cells per trajectory ranged between 10 and 60 cells.

To confirm the observed negative correlation between start ratio and maximum response, we performed a similar classification on the data from the first experiment, shown in Figure 14. As observed in Figure 21, we found again that the amplitude of the responses is highest among the cells with the lowest start ratios.

**Figure 21.**
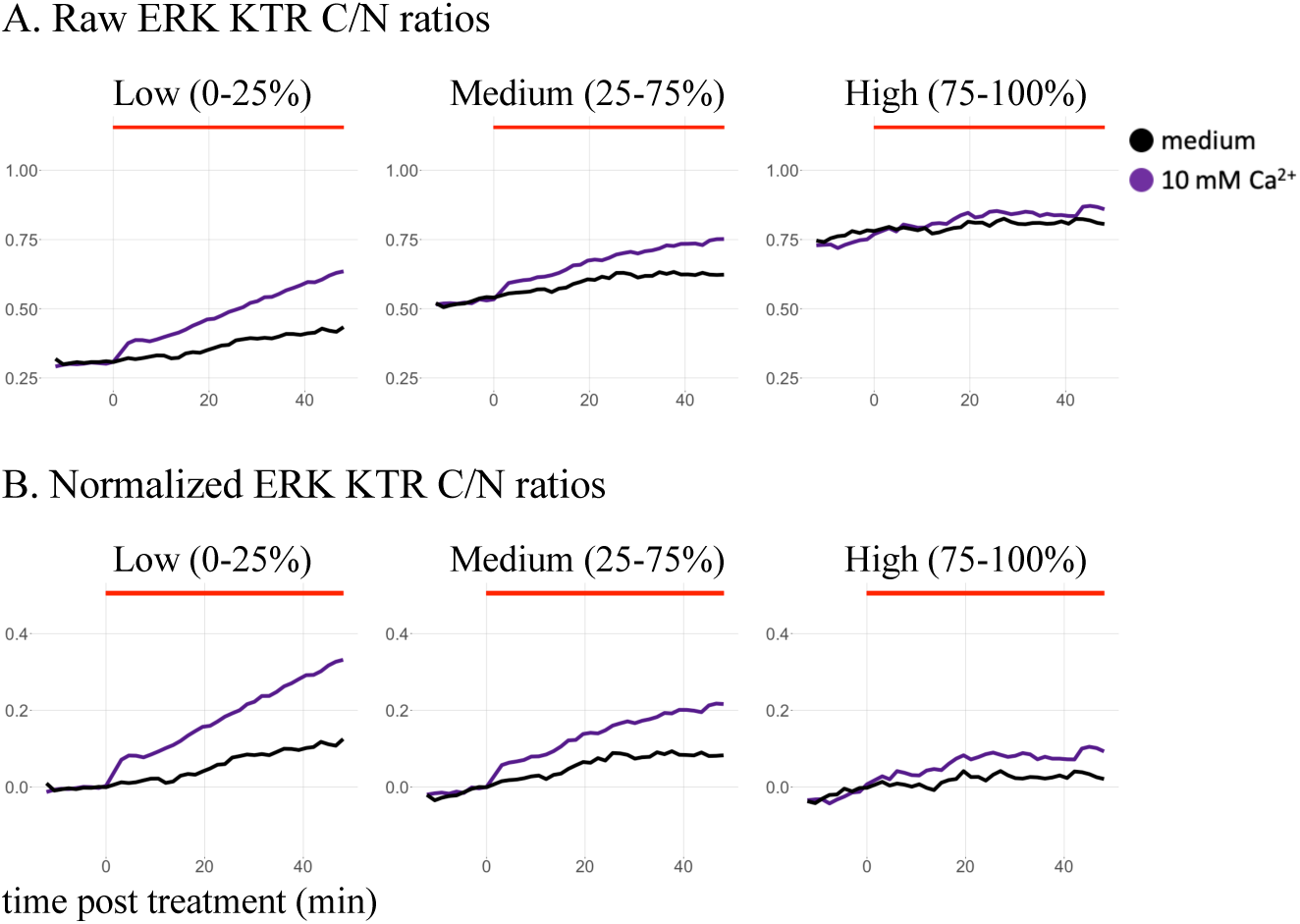
Effect of start activity over ERK responses over time upon calcium stimulation. A: Raw data. B: Normalized data to C/N ratio prior to stimulation. Cells were pre-treated with vehicle DMSO or with 1 μM NPS-2143, and stimulated with medium or 10 mM Ca^2+^. Red bar indicates stimulation. Black and purple colored trajectories correspond to treatments with medium and 10 mM Ca^2+^ respectively. Dots indicate the mean and thin lines show the standard deviation. The numbers of cells per trajectory ranged between 10 and 60 cells.

These data altogether suggest that calcium stimulation results in ERK activation in a concentration-dependent manner, and that pre-treatment with CaSR inhibitor limits these effects.

## DISCUSSION

Here, we present our attempt at combining two different biosensor technologies to study simultaneously in single cells the behaviour of an early and a downstream component within the CaSR signaling pathways. This novel approach incorporating previously validated Gi protein FRET-based and ERK translocation biosensors turned out to be very technically challenging. Despite substantial efforts to optimize the generation of the cell line, image processing, segmentation and tracking, and data analysis, obtaining robust single-cell data for both biosensors was unsuccessful.

We generated a stable cell line expressing two sensors and a nuclear marker to study the Gi-ERK pathway in single HEK-CaSR cells. The constructs were expressed as expected, with the Gi sensor locating predominantly in the membrane, the H2A marker in the nucleus, and the ERK KTR translocating between the nucleus and the cytoplasm. Through extensive fluorophore characterization and image processing, we generated a pipeline to unmix the individual fluorescent protein signals in the cells. Examination of the CFP and YFP counts over time in single cells showed similarities with those from GFP and OFP, suggesting insufficient unmixing. Considering the magnitude of expected changes in CFP and YFP counts upon activation of the FRET sensor, the effects of an imprecise unmixing from the translocating GFP interfered with the quantification of YFP and CFP intensities. As a consequence, it was not possible to obtain reliable YFP/CFP ratios. In terms of ERK, we found indications of ERK activity in response to calcium stimulation, possibly via CaSR. These observations were especially clear after classifying the trajectories based on the start ratio prior to stimulation, which had a major effect on the measured changes. The present study shows the challenges of combining these two technologies, and at the same time points to potential improvements to overcome these limitations.

### Generation of stable cell line

In this project we have used the PiggyBac transposon system to generate stable cell lines that express two different constructs. The integration efficiency for single transfections was very high, with puromycin generating up to 98% fluorescence-positive cells after 2-5 days antibiotic selection, and blasticidin up to 70% after 7 days (data not shown). However, the efficiency decreased tremendously with the simultaneous double transfection, limiting the availability of monoclonal populations with the desired expression levels of both constructs. This low efficiency could be partially explained by the transposase and the transposon vector. The used transposase has both insertion and excision activities, decreasing the chances of simultaneous integration of two constructs. In addition, the transposon vector had two different promoters, one for the antibiotic resistance, and one for the construct of interest, allowing for single expression of the antibiotic resistance gene under certain conditions.

Using an alternative transposon system, or lentiviral transduction [31], could increase the efficiency of integration of multiple constructs and thus result in a larger pool of monoclonal populations. These populations could be further compared and tested, as performed in an earlier study [18], to find a better balance in the expression of both plasmids and potentially measure simultaneously changes in FRET and ERK.

### Probe selection

For this project, we wanted to use single excitation for all fluorophores, as this increases imaging speed, and reduces optical artifacts. This approach limits the choice of the 3rd and 4th fluorophores to large-Stokes-shift green and orange/red fluorescent proteins. From the available lssGFPs, we selected lssSGFP2, engineered in our lab, as it outperformed others in brightness, tendency to oligomerization, and photostability [24]. Out of the tested lssOFPs/RFPs, lssmOrange was brighter than LssmCherry1, LssmKate2 and hmKeima8.5 in single cells. Due to the higher dimerization tendency of lssmOrange compared to lssSGFP2 [24], we decided to use lssSGFP2 for the KTR, to minimize any unknown impact of dimerization over the performance of the KTR. However, a recent study has shown that oligomerization could increase the dynamic range of KTRs, possibly by increasing its size and minimizing its passive diffusion through the nuclear pore [32]. In addition, in the same study, the authors optimized the NLS sequence in the ERK-KTR to generate a variant with improved nuclear localization and dynamic range in HEK-293 cells. Unsurprisingly, a different variant yielded better results in neurons, highlighting the potential of optimizing the KTR design for the cell type of interest to account for intrinsic characteristics in terms of nuclear import/export dynamics and basal kinase activities.

Optimizing the NLS sequence and tagging the ERK-KTR with the less monomeric lssmOrange could potentially increase its dynamic range, and at the same time facilitate the measurement of FRET changes by using lssSGFP2 to track the non-dynamic nuclear marker. Another potential advantage of swapping these fluorescent proteins, is that lssSGFP2 is more photostable than lssmOrange [24]. A stable nuclear marker signal over time facilitates precise nuclear segmentation, while the consequences of partial bleaching in a KTR are limited as they would affect both compartments.

Alternatively, considering the implications of the high bleedthrough from lssmOrange into the YFP channel, lssRFP hmKeima8.5 could have been a better choice as it would not be detected in that channel. Recently published lssRFPs mCRISPRed and lssmScarlet3 [33,34], with a theoretical brightness similar to hmKeima8.5, could also be suitable candidates.

### Image processing

The imaging strategy used here required a great effort in image processing and analysis. Several steps and controls were necessary to obtain aligned, shading-corrected, and bleedthrough-corrected images.

During acquisition of the images, we noticed a minor misalignment between the 4 quadrants, which was resolved using a plugin that aligns the images based on the overlap of common structures. High bleedthrough and co-localization of the fluorophores facilitated the alignment of the GFP, YFP and OFP channels, while the alignment of membrane-localizing CFP was occasionally imprecise when the object density in the field of view was too low.

During the calculation of the bleedthrough coefficients in single transfected cells, we found differences of up to 15% between individual measurements. Part of this variability could be attributed to shading, which resulted in differences of up to 10% in the same cell upon correction. Despite the clear effects in coefficients in individual cells, shading correction had no effect at the population level, yielding comparable average coefficients but with a lower spread in the data distribution.

Given the observed spread in the measured coefficients between individual cells, it was expected that using a calculated average could result in under- or over-segmentation. This was directly observed among cells in the YFP channel, where the unmixing resulted in negative values and the coefficients had to be reduced. Imprecise segmentation affects directly the obtained measurements, but the consequences on the data depend on multiple factors. If the biosensor co-localizes with a stable static fluorophore, such as a nuclear marker, removing too many or too few counts from the biosensor channel will affect the absolute counts and possibly the amplitude of the measured changes as well. This effect can be considerable for FRET biosensors read-outs, where the expected intensity changes of the individual fluorophores are small. On the other hand, the effect on the C/N KTR ratios will be less severe as the intensity changes are much larger in comparison. In our case, where the FRET sensor also co-localizes with the KTR, the large changes in intensity resulting from the translocating KTR drastically affects the FRET measurements. Expectedly, our measurements in single cells showed similarities in the relative changes of the counts over time in all 4 channels, suggesting an imperfect segmentation. Nevertheless, we could observe and measure clear KTR translocations, as a combined result of the high intensity counts of the KTR, the limited bleedthrough from other channels, and the very high dynamic range. Unfortunately, for the opposite reasons, we were not able to measure changes in the FRET sensor. This was the case not only in the nucleus but also in the cytoplasm, suggesting that the large bleedthrough from the dynamic lssSGFP2 into the CFP and YFP channels alone is sufficient to interfere with the FRET measurements.

Using lssSGFP2 to tag the non-dynamic nuclear marker, and tagging the mobile ERK-KTR with lssmOrange or a lssRFP, would minimize the bleedthrough into the CFP and YFP channels and possibly be sufficient to obtain robust FRET ratio changes. Yet, the FRET biosensor by itself shows relatively low fluorescence. The reason is that heterotrimeric G-protein FRET sensors are typically expressed at low levels to minimize interference with normal cell physiology. Moreover, low expression is often required to get a good response. Since the photons emitted by the YFP are from indirect YFP excitation and the FRET process, the signal in this channel is much lower than for directly excited YFP. Altogether, it is hard to improve the brightness of the FRET based biosensor, as the brightest available FPs are used. A more realistic improvement would be to increase the dynamic range, but this would still require substantial engineering of the biosensor. Employing a tandem acceptor, as in Epac-based sensors [35], would be the simplest change to try.

### Segmentation and tracking

Successful segmentation and tracking of the individual nuclei and cytoplasms over time is fundamental to extract accurate information from the images. Generation of nuclear masks with StarDist worked well in our data, as shown in Figure 9. However, StarDist uses star-convex polygons and it is known to work best with round objects. Most of the imaged nuclei were round or oval, although a fraction of them were mildly kidney shaped, which led to single nuclei being considered as two adjacent nuclei and to the edges being smoothened to be rounder. Splitting of nuclei in some of the frames reduces the number of single tracked objects over the timelapse, and smoothening of the edges affects the accuracy of the mask. A later published segmentation tool, CellPose [36], could produce more accurate masks as suggested by a recent study comparing both tools in a similar dataset [37]. Additionally, further improvements could be done in the segmentation of the cytoplasms, which were identified by expanding the segmented nuclei using a cellular binary mask generated by blurring a thresholded image that combined the counts from the KTR and H2A channels counts. This approach worked very well overall, but establishing the boundary between background and foreground can be imprecise when the cytoplasmic KTR signal is very low due to strong nuclear localization. An option to address this is to use the CFP channel images to identify the cell membranes, with tools such as CellPose or DeepCell [38], which have shown to outperform other segmentation tools with membrane labeled cells [37].

To follow the behavior of the single cells over time, we used the overlap tracking method from CellProfiler. It is a rather simple but effective method to use when there is a high spatial overlap of the objects between frames. Despite the mobility of the cells, the necessary overlap was achieved by intentionally imaging fields of view with a low cell density and using the minimum possible pixel distance.

The use of these novel segmentation algorithms could improve the accuracy of the nuclear and cellular segmentation and thus greatly increase the number of tracked objects over time, fundamental for single cell heterogeneity analysis.

### CaSR activation and ERK signaling

Our experimental set-up allowed to track the ERK KTR C/N ratios over time in HEK-293 cells stably expressing CaSR. Activation of ERK by CaSR has been well established in various cell types and tissues, via pathways involving Gq, Gi, and β-arrestins [5]. Studies in HEK-293 cells expressing CaSR have shown that ERK phosphorylation reaches a maximum about 10 min after calcium stimulation and is still visible after 60 min [11,39]. In addition, a study using tetracycline-inducible CaSR in HEK-293 cells showed that negative allosteric modulators such as NPS-2143 reduce both the potency and the maximum response of calcium to activate ERK, but the effect is greater for intracellular calcium mobilization [40].

Our previous work on KTRs to study activation of endogenous GPCRs [18] showed that ERK-KTR has a large dynamic range and is a powerful tool to study cell-to-cell heterogeneity as it can be used to track ERK activity in hundreds of cells with relatively good temporal resolution. Of note, that study was done in HeLa cells, which are less mobile and show little morphological responses to GPCR activation. As such, the careful selection of a cell line that is used for tracking the response to CaSR activation with imaging methods is recommended.

Our results showed higher average rates of translocation upon stimulation with increasing concentrations of calcium, as well as decreased translocation in the presence of NPS-2143. However, the observed differences were modest, and the average trajectories did not reflect clearly the kinetics reported in literature. In addition, we found a negative correlation between start ratio and maximum response, as well as slow KTR translocation over time upon medium addition. These factors, combined with the high heterogeneity between cells within individual fields of view in terms of start ratios and kinetics over time, limited the comparison between the different conditions and therefore the interpretation of the data.

Due to the aforementioned limitations, we were not able to generate large amounts of robust single cell data for Gi protein and ERK activation, and we decided to not pursue this line of investigation. However, the observed single cell ERK responses and heterogeneity remain interesting phenomena and should be further explored. For this purpose, we have included various suggestions to optimize the transduction, biosensor design, and segmentation. Nevertheless, there are several questions that could be answered with the current model.

To confirm the role of CaSR in the measured ERK responses to different concentrations of extracellular calcium, we could make use of CaSR positive allosteric modulator NPS-R568 and a combination of Gq and Gi protein inhibitors, which should increase and decrease CaSR activity respectively.

Aside from CaSR signaling, and despite the clear limitations we found, this model might be of use to better understand the potential usability of ERK-KTRs in HEK-293 cells. To the best of our knowledge, only two other studies have utilized any KTR in HEK-293 cells. One study is the original publication describing KTRs, where the authors showed JNK-KTR translocation upon treatment with Anisomycin [17], and the other study evaluated various engineered variants of ERK-KTR in response to EGF in HEK-293 cells and neurons [32]. Attempting to replicate these results would be a first step, as well as stimulating endogenously expressed GPCRs such as LPARs or S1PRs [41] and comparing these responses to available ERK phosphorylation data. For these experiments, normal extracellular calcium concentrations in the medium would be preferred as they would facilitate attachment of the cells. These additional data can also help corroborate and understand the negative correlation between the start KTR ratio and the maximum measured response, which we observed in this study but not in HeLa cells in a previous study [18]. Moreover, stimulating these GPCRs could also be used to confirm whether the absence of clear FRET ratio changes was due to the ERK-KTR translocation and imprecise unmixing of the signals. Pre-incubating the cells with an inhibitor upstream of ERK, such as PD 0325901, and imaging the cells every 1-2 s should be sufficient to observe clear FRET ratio changes upon Gi activation.

Last, it will be important to investigate the observed slow continuous ERK translocation overtime in the absence of stimulation. A possible explanation is that the mix of MM and FluoroBrite had an insufficient HEPES concentration to prevent an increase in pH in the medium while imaging at 37 °C without CO_2_, and this led to alkalosis-dependent ERK phosphorylation [42]. If the pH of the medium does increase over time in the absence of COand leads to ERK translocation, increasing the HEPES concentration in the medium could easily solve this.

Finally, if efforts to effectively combine a FRET-pair and a KTR prove insufficient or too complex, a single excitation imaging system could still be used to quantify 3 KTRs and a nuclear marker, using a CFP, a lssGFP, a lssYFP, and a lssRFP/OFP. Theoretically, two FRET-sensors could also be measured simultaneously, including a CFP-GFP/YFP pair and a lssGFP/lssYFP-RFP pair. However, this approach is less straight-forward due to the low availability of optimized lssGFP/lssYFP-RFP FRET-based biosensors.

## METHODS

### Reagents

CaCl_2_ dihydrate (Sigma-Aldrich, # 21097) was prepared as a 1 M solution in ddH2O. NPS-2143 hydrochloride (Tocris, # 3626) was prepared as a 1 mM solution in DMSO.

### Cloning

To generate the tandem constructs containing each FP and mTq2 with a T2A linker in between, we replaced mNG from mNG-T2A-mTurquoise2 with LssmCherry1, LssmKate2, hmKeima8.5 or LssmOrange. First, mNG-T2A-mTq2 and the FP of interest, in a Clontech-style C1 plasmid, were cut using the restriction enzymes NdeI and Kpn2I. The digested products were then ligated using T4 DNA ligase, and the products were transformed by heat-shock in DH5a *Escherichia coli* competent cells.

To clone the G-protein FRET-based biosensors into PiggyBac transposon vectors containing a puromycin resistance gene [7–9], the biosensors for Gαi1, Gαi2 and Gαi3 were first amplified by PCR (Fw: 5’-TATA<u>GAATTC</u>CCGTCAGATCCGCTAG-3’ and Rv: 5’-TATA<u>GAATTC</u>ATGTGGTATGGCTGATTATGATC-3’) to introduce EcoRI restriction sites (underlined in primers sequences) upstream and downstream of the biosensor. The Gαq and Gα13 sensors were amplified by PCR (Fw: 5’-TATA<u>ACTAGT</u>CCCCGTCAGATCCGCTAGC-3’ and Rv: 5’-TATA<u>GAATTC</u>ATGTGGTATGGCTGATTATGATC-3’) to insert SpeI and EcoRI sites upstream and downstream sites (underlined in primers sequences). For the Gαq sensor, we first introduced a silent mutation using SDM by PCR (Fw: 5’-GCAGCCCGAGA<u>G</u>TTCATTCTGAAGATGTTCG-3’ and Rv: 5’-CAGAATGAA<u>C</u>TCTCGGGCTGCCTGGG-3’) to remove an EcoRI restriction site located within the α subunit (mutation underlined in primers sequences). For this, we used PfuTurbo DNA polymerase, followed by DpnI digestion to destroy template DNA.

To clone the sensors into the piggyBac transposon vector pMP-PB [43], kindly shared by Jakobus van Unen and David Hacker, we digested the G-protein amplicons and the vector with the mentioned restriction enzymes and ligated them using T4 DNA ligase, and transformed the products by electroporation using *Escherichia cloni* competent cells.

To generate the construct containing the ERK-KTR and a nuclear marker, we first tagged the P2A-ERK-KTR and the H2A sequences with lssSGFP2 (Mastop et al. unpublished data) and lssmOrange respectively [22], using AgeI and BsrGI to replace the fluorescent proteins from H2A-mScI and P2A-ERK-KTR-mNG, described in chapter X. The P2A sequence was kept to ensure equimolar expression of the separate proteins from a single transcript [23]. Then, we ligated P2A-ERK-KTR-lssSGFP2 digested with Acc65I and BsrGI and H2A-lssmO digested with BsrGI, to generate H2A-lssmO-P2A-ERK-KTR-lssSGFP2, which we refer to as “HOEG”. This ligation destroys the Acc65I/BsrGI site in between the KTR and the H2A. Afterwards, we cloned HOEG into the piggyBac transposon vector pMP-PB containing a blasticidin resistance marker using EcoRI and NotI. All ligation products were transformed using electro competent *E. cloni* cells.

### Fluorescent protein isolation

For protein purification of mTq2, lssSGFP2, cpVenus and lssmOrange, we first transformed electro competent *E. cloni* cells with pDX vectors containing each FP. These vectors allow for protein expression in bacteria under a rhamnose promoter, and in mammalian cells, under a CMV promoter.

Transformed bacteria colonies were grown overnight at 37 °C with agitation (200 rpm) in 50 ml super optimal broth (SOB, 0.5% (w/v) yeast extract, 2% (w/v) tryptone, 10 mM NaCl, 20 mM MgSO4, 2.5 mM KCl in ddH2O) supplemented with 0.4 % (w/v) L-rhamnose (Sigma-Aldrich, cat#R3875) and 50 μg/ml kanamycin, and for additional 6 h at 20 °C L (to promote protein synthesis or to improve maturation). Bacteria were then spun down for 30 min at 3220 g, and the pellet was resuspended in 20 ml salt-Tris buffer (ST, 20 mM Tris-HCl, 200 mM NaCl in ddH2O, pH=8.0). The suspension was spun down for 30 min at 3220 g at 4°C, and the pellet was resuspended in 5 ml ST supplemented with 1 mg/ml lysozyme (Sigma-Aldrich, cat#L7651) and 5 U/ml benzonase nuclease (Millipore, cat#71205-3) and incubated on ice for 30 min. Then, we added 100 µl 100 mM PMSF and 100 µl 10% NP40, sonicated for 5 min at 40W, and spun down for 30 min at 40 000 g at 4°C. The supernatant was added to 1 mL Ni-NTA His-Bind resin (Novagen, cat#69670-2) and incubated at 4°C for 1 hour with rotation. The resin was then washed three times with ST buffer by spinning down for 5 min at 550 g. The resin was incubated for 10 min in 0.5 ml 0.6 M imidazole in ST, and spun down for 5 min at 550 g to collect the supernatant. The resin was eluted a second time with 0.5ml 0.2 M imidazole in ST, and the collected supernatant mixed with the first supernatant. The supernatant was then sterilized with a 0.22 μm filter, and dyalized overnight in 2L 10 mM Tris pH=8 at 4°C using a 3.5 kD membrane tubing (Spectrum Laboratories, cat# 132720).

### Cell culture and transient transfection

HEK-293 and HeLa cells were obtained from the American Tissue Culture Collection (ATTC, Manassas, VA, USA), and HEK-CaSR cells were a kind gift from Dr. Donald Ward, University of Manchester. HeLa and HEK-CaSR cells were grown in full growth medium, Dulbecco’s Modified Eagle Medium with GlutaMAX (Gibco, cat# 10566016) supplemented with 10% fetal bovine serum (FBS) (Gibco, cat# 10270106), at 37°C in 7% COin humidifying conditions. Cells were passaged every 2-3 days by washing with HBSS (Gibco, cat# 14170), trypsinizing using 0.25% Trypsin-EDTA (Gibco, cat# 25200056), spinning down at 300 g for 5 min, and resuspending in full growth medium. All cells were routinely tested for mycoplasma by PCR.

For transient transfection, 300 000 −400 000 HEK-293 or 400 000 −500 000 HeLa cells were seeded two days in advance on a 35 mm diameter well containing a 24 mm diameter glass coverslip (Thermo Scientific Menzel, cat# 11778691). One day before the experiment, the cells were transiently transfected with the constructs of interest using Polyethylenimine (PEI) (1 mg/mL in ddH2O). 0.5 μg of DNA per plasmid was mixed with PEI at a ratio 1:4.5 in 100 μL OptiMEM (Gibco, cat# 31985).

### Spectral imaging microscopy

We acquired spectral images of single HeLa cells transiently transfected with tandem constructs of mTq2 and a lssOFP or lssRFP with a T2A sequence in between, as previously described [44]. Spectral images were obtained by exciting the cells with 436/20 nm, an 80/20 transmission/reflection dichroic, and a 460 LP emission filter. Cellular ROIs were drawn manually and the spectral images were saved as “ics” files and subsequently extracted in FiJi with the function “Plot profile”. The spectra were then normalized to the mTq2 emission peak at ∼505 nm, and the mTq2 spectrum, obtained from cells transfected with mTq2 alone, was subtracted to obtain the emission spectra of lssOFP/lssRFP relative to mTq2.

### Quantification of brightness in cells

HeLa cells were transiently transfected with tandem constructs of mTq2 and several lssOFP/RFPs with a T2A sequence in between. Cells were imaged using a wide-field fluorescence microscope (Axiovert 200M, Carl Zeiss GmbH) kept at 37 °C, equipped with an oil immersion 40x objective (Plan-Neo-fluor 40× /1.30; Carl Zeiss GmbH). The fluorescent proteins were illuminated using a xenon arc lamp (Cairn Research, Faversham, Kent, UK) with a computer-controlled monochromator. The images were binned 4×4 and recorded with a cooled charged-coupled device camera (Coolsnap HQ, Roper Scientific, Tucson, AZ, USA), controlled with software MetaMorph 6.1 (Molecular Devices). CFP, lssOFP (lssmOrange), and lssRFP (hmKeima8.5, lssmKate2, and lssmCherry1) channels were measured by exciting the sample with 450 nm, 470 nm, and 480 nm light respectively, reflecting the light using a 455, 552, and 585 dichroic mirrors, and detecting consecutively emission with a 470/30, 579/34, and 620/60 bandpass filters.

Exposure time was 100 ms for all channels, and 5 sequential images were acquired per channel. After subtracting the background, the average intensities of the 5 images per cell per channel were calculated, and the lssOFP/lssRFP intensities were corrected for bleedthrough from mTq2, which was calculated to be 40% and 5% respectively. Finally, the fluorescence intensities of each lssOFP/lssRFP were plotted against mTq2 in the same cell to represent relative brightness [45].

### Live-cell imaging

For live-cell imaging, we used a Nikon Eclipse Ti-E inverted wide-field microscope (Nikon) equipped with a 20X and 40X objectives (Plan Apo 20X NA 0.75 Air, Plan Apo 40X NA 0.95 Air) and an Intensilight mercury lamp (C-HGFIE, Nikon). The microscope stage and filter wheels were controlled by a ProScan III unit (ProScan H31, Prior Scientific) through the Nikon NIS elements AR 4.10.

To acquire the images, we excited the cells with 420/40 nm (AT420/40, Chroma) and a 450nm dichroic mirror (ZT442rdc, Chroma). The emission light is first passed through a HQ460 LP filter (Chroma) to reflect residual excitation light, and detected by an ORCA-Flash 4.0 V2 digital CMOS camera (C11440-22CU, Hamamatsu Photonics) in combination with a Cairn Multisplit V1. The Multisplit unit contains three dichroic filters, a primary of 525 nm, and two secondaries of 495 and 560 nm, which results in 4 channels: 460-495 nm, 495-525 nm, 525-560 nm, and >560 nm. Exposure times ranged from 500 to 800 ms, and camera binning was set to 2×2.

### Determination of bleedthrough coefficients

The experimental determination of bleedthrough coefficients was performed in transformed bacteria, beads with purified protein, and transfected HEK-293 cells.

Electro competent *E. cloni* cells were transformed with pDX vectors containing mTq2, lssSGFP2, cpVenus and lssmOrange, as well as with a plasmid without the pDX promoter as control. Bacteria were plated on a Petri dish with super optimal broth agar (SOB, 0.5% (w/v) yeast extract, 2% (w/v) tryptone, 1.5% (w/v) agar, 10 mM NaCl, 20 mM MgSO4, 2.5 mM KCl) supplemented with 0.4 % (w/v) L-rhamnose (Sigma-Aldrich, cat#R3875) and grown overnight at 37°C. The following day, the plates were further incubated at 4°C for 96 hours for complete maturation of the protein.

Nickel NTA beads (Novagen) were incubated with purified mTq2, lssSGFP2, cpVenus and lssmOrange for 1 hour at 4°C, washed with ST buffer (20mM Tris 200mM NaCl in ddH2O, pH=8).

HEK-293 cells were transiently transfected overnight with OptiMEM, PEI, and a single construct containing mTq2, lssSGFP2, cpVenus or lssmOrange, either targeted to the nucleus or non-targeted.

Subsequently, the bacterial colonies, beads, and HEK-293 cells were imaged as mentioned above using the Multisplit system. From the images, we manually drew ROIs in the bacterial colonies, beads, and single cells, as well as in background areas and the control transformed bacteria. We then extracted the mean intensity values of these ROIs and subtracted the corresponding background. After obtaining the mean value per channel, the values were normalized to the channel of the measured fluorophore.

### Generation of stable cell lines

To generate the stable cell lines, we used the NeonKit (ThermoFischer) to electroporate HEK-CaSR cells with transposase and the transposon vectors PB_Blasticidin_HOEG and PB_Puromycin_Gi1-mTq2-cpV.

We first trypsinized the cells, and resuspended them in buffer R at 14×10^6^ cells/mL density. 4.5 μg of transposon vector and 3 μg of transposase were added to 5 million cells, and the mix was subjected to 2 pulses of 20 ms at 1100 V. The electroporated cells were plated in full growth medium, expanded to reach 60-70% confluency, and selected for 96 h with 1 μg/mL Puromycin (Gibco, cat# A1113803) and 2.5 μg/mL Blasticidin. The cells were passaged to reach confluency in T25 flasks.

### FACS

To sort by fluorescence-activated cell sorting (FACS), the cells were first washed, trypsinized, spun down, and resuspended in full growth medium as for passaging. Then, the cells were spun down, resuspended in 2% FBS in HBSS containing 1 μg/mL DAPI (Invitrogen, cat# D1306), spun down, resuspended in DAPI-free 2% FBS-HBSS, and kept in the dark on ice. Cells were sorted with the FACSAria™ III (BD Biosciences, Franklin Lakes, NJ, USA), using a 100 μm nozzle at 20 psi pressure.

Single cells were identified by drawing gates using the area, width, and height of forward scatter (FSC) and side scatter (SSC), and living cells based on being DAPI-negative. Living cells were identified based on the DAPI staining. Due to the spectral overlap of the fluorophores, we used samples with cells expressing the individual fluorophores, or fluorophore pairs, to unmix the different signals. DAPI was excited with 405 nm and measured with a 450/50 nm bandpass emission filter. cpVenus and lssSGFP2 were excited with 488 and 405 nm respectively, and detected with a 530/30 and 510/50 bandpass emission filters. First, we set two gates based on cpVenus fluorescence for “medium” and “high” expression of the FRET pair. Then, we used the lssSGFP2 intensity to obtain two gates for “low” and “medium” expression of the H2A-KTR pair.

We then sorted the pools of the different intensity combinations into 15mL tubes. The tubes and plates contained full growth medium, supplemented with 10 mM HEPES and 1% Penicillin/Streptomycin (P/S) (Gibco, cat# 15140148). From these pools, single cells were obtained by serial dilution in fibronectin-coated 96-well plates, and sequentially transferred to bigger wells/flasks to expand.

### Calcium stimulation experiments

We used Ibidi 8-well glass bottom slides (Ibidi, cat# 80827) and coated them with 35 μg/ml of poly-D-lysine (Sigma-Aldrich, cat#P6407). Unless stated otherwise, 8-24 h prior to the experiment 200 000 cells were seeded per well in full growth medium, and 2 h before imaging the medium was removed and replaced with a 1:1 mix of Fluorobrite (Gibco, cat# A1896701) and 0 mM Ca^2+^ microscopy medium (MM) (20 mM HEPES pH=7.4, 137 mM NaCl, 5.4 mM KCl, 0.8 mM MgCl2, 20 mM Glucose) to reduce the basal kinase activity of ERK. During imaging, the cells were pre-treated with vehicle (0.033% DMSO) or 1 μM CaSR inhibitor NPS-2143, and treated with medium (1:1 Fluorobrite:MM) or 3, 5, or 10 μM Ca^2+^.

### Image processing

Unless stated otherwise, image processing was performed using FIJI [46]. First, the images were split into 4 quadrants of equal size, and to correct for the imperfect alignment of the dichroics, the quadrants were aligned by translation using the plugin “StackReg” [47]. Second, the quadrants were corrected for uneven illumination and detection (i.e. shading), for which we used the following formula:

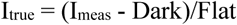

I_meas_ represents the measured counts, and I_true_ shading-corrected counts. To calculate “Dark”, we first generated the average image of a time-lapse (200 images 500 ms binning 2×2) without any light source. After removing the bright counts using “Remove outliers” with a radius of 1 pixel, we calculated the average pixel intensity per quadrant. This value, “Dark”, ranged between 400 and 430 counts. To calculate “Flat”, flat-field images were obtained by imaging fluorescent protein solutions. Per quadrant, after subtracting the dark counts, we normalized the counts to the quadrant average, and applied a Gaussian blur of radius of 15 pixels. For each FP, “Flat” corresponded to the quadrant with the highest counts.

Third, the 4 signals were linearly unmixed by applying using an unmixing matrix MATLAB script and an unmixing matrix. Finally, the background was removed by subtracting the average intensity of a manually drawn ROI without cells. These steps resulted in the processed images that were used for quantification of fluorescence intensity or further processed as indicated below for segmentation purposes.

Whenever mentioned, the tool “Subtract background” was also tested and used, with a rolling ball of 55 pixels radius for the RFP channel or 90 pixels radius for the other channels.

To facilitate nuclear segmentation, we applied the tool “Subtracted background” to the processed RFP images as mentioned above, then subtracted 5 000 counts, and applied a Gaussian Blur with sigma radius 1.

To identify the area covered by cells in a field of view, we first applied the “Subtracted background” tool to the processed GFP and RFP images as specified above. Then, we mathematically added the counts of both images, applied the “Auto threshold” tool with the Huang2 method, and applied a Gaussian blur with sigma radius 2.5.

### Segmentation and tracking of nuclei and cytoplasms

To segment the nuclei, we applied the StarDist [26] plugin in ImageJ (default settings) to the previously processed nuclear images. Then, in CellProfiler (version 4.1.3) [27],we used the segmented nuclear objects (i.e. regions of interest or ROI) as seeds to identify the cellular ROIs. For this, we used the previously generated images that combined the GFP and RFP counts, and nuclei masks were expanded by 15 pixels in all directions as long as there was no background. The first expanded pixel was discarded to ensure clear separations between nucleus and cytoplasm. The nuclear and cellular ROIs were then tracked through the time lapses based on overlap. These ROIs were identified as unique objects if the distance between their positions in consecutive images was lower or equal to 5 pixels. Finally, the size/shape features of the nuclear and cytoplasmic ROIs were exported, together with the intensity features of these ROIs in the background-subtracted mTq2, lssSGFP2 and cpV images.

### Data processing and plotting

We used a custom-made R [48] script to process the exported data from CellProfiler and to generate the plots. First, we filtered the data to remove objects with suboptimal segmentation or tracking. For this purpose, we excluded cells with nuclear/cytoplasmic areas lower than 200 pixels, and with mean intensity values lower than 1500, 1000, and 500 counts for GFP, CFP, and YFP. In addition, we removed the objects that were not present in each time point as a single object, objects that were too distant between consecutive frames (10 or more pixels), and objects with short trajectories (shorter than the maximum minus 7 frames). Then, we calculated the ERK CN ratios for the lssSGFP2 channel by dividing the mean intensities of the cytoplasmic by the nuclear ROIs, and the Gi FRET ratios by dividing the mean intensities of the YFP by the CFP. For normalization, we used the average of the two last measurements prior to stimulation, and the FRET and CN ratios were normalized by division and subtraction, respectively.

